# Non-indigenous seaweeds in the Northeast Atlantic Ocean, the Mediterranean Sea and Macaronesia: a critical synthesis of diversity, spatial and temporal patterns

**DOI:** 10.1101/2023.06.05.543185

**Authors:** Luna M. van der Loos, Quinten Bafort, Samuel Bosch, Enric Ballesteros, Ignacio Bárbara, Estibaliz Bercibar, Aurélie Blanfuné, Kenny Bogaert, Silke Bouckenooghe, Charles-François Boudouresque, Juliet Brodie, Ester Cecere, Pilar Díaz-Tapia, Aschwin H. Engelen, Karl Gunnarson, Soha Hamdy Shabaka, Razy Hoffman, Vivian Husa, Álvaro Israel, Mart Karremans, Jessica Knoop, Line Le Gall, Christine A. Maggs, Frédéric Mineur, Manuela Parente, Frank Perk, Antonella Petrocelli, Conxi Rodríguez-Prieto, Sandrine Ruitton, Marta Sansón, Ester A. Serrão, Adriano Sfriso, Kjersti Sjøtun, Valérie Stiger-Pouvreau, Gwladys Surget, Thierry Thibaut, Konstantinos Tsiamis, Lotte Van De Weghe, Marc Verlaque, Frédérique Viard, Sofie Vranken, Frederik Leliaert, Olivier De Clerck

## Abstract

Effective monitoring and combatting the effect of non-indigenous seaweeds relies on a solid confirmation of the non-indigenous status of the species. We critically analysed the status of presumed non-indigenous seaweed species reported from the Mediterranean Sea, the Northeast Atlantic Ocean and Macaronesia, resulting in a list of 140 species whose non-indigenous nature is undisputed. For an additional 87 species it is unclear if they are native or non-indigenous (cryptogenic species) or their identity requires confirmation (data deficient species). We discuss the factors underlying both taxonomic and biogeographic uncertainties and outline recommendations to reduce uncertainty about the non-indigenous status of seaweeds. Our dataset consisted of over 19,000 distribution records, half of which can be attributed to only five species (*Sargassum muticum*, *Bonnemaisonia hamifera*, *Asparagopsis armata*, *Caulerpa cylindracea* and *Colpomenia peregrina*), while 56 species (40%) are recorded no more than once or twice. In addition, our analyses revealed considerable variation in the diversity of non-indigenous species between the geographic regions. The Eastern Mediterranean Sea is home to the largest fraction of non-indigenous seaweed species, the majority of which have a Red Sea or Indo-Pacific origin and have entered the Mediterranean Sea mostly via the Suez Canal. Non-indigenous seaweeds with native ranges situated in the Northwest Pacific make up a large fraction of the total in the Western Mediterranean Sea, Lusitania and Northern Europe, followed by non-indigenous species with a presumed Australasian origin. Uncertainty remains, however, regarding the native range of a substantial fraction of non-indigenous seaweeds in the study area. In so far as analyses of first detections can serve as a proxy for the introduction rate of non-indigenous seaweeds, these do not reveal a decrease in the introduction rate, indicating that the current measures and policies are insufficient to battle the introduction and spread of non-indigenous species in the study area.

**Highlights:**

- Non-indigenous seaweed species in the Northeast Atlantic Ocean, the Mediterranean Sea and Macaronesia are critically reanalysed.
- >19,000 distribution records revealed considerable variation in diversity of non-indigenous seaweed species in the study area.
- Taxonomic and biogeographic uncertainties hamper a critical evaluation of the non-indigenous status of many seaweed species.

## Introduction

Over the course of several centuries, human-mediated transport has led to the introduction and establishment of more than 14,000 non-indigenous species in Europe (EASIN, 2022). Some of these non-indigenous species profoundly affect the abundance, diversity, interactions and evolution of native biota and consequently affect ecosystem structure, functions and services (Simberloff *et al*., 2013; Dawson *et al*., 2017; Blakeslee *et al*., 2020). The introduction of non-indigenous species can also result in substantial negative economic impacts (Hulme *et al*., 2009). The reported costs of biological invasions, at a global level, were estimated to be at least 1.288 trillion US Dollars over 1970–2017 (Diagne *et al*., 2021). Furthermore, biotic homogenisation and consequently also the impact of non-indigenous species on native ecosystems are expected to increase in the context of climate change (Bennett *et al*., 2021).

The management of biological invasions depends heavily on lists of reliably identified non-indigenous species. Such lists form an essential tool underpinning prevention, control, mitigation or eradication strategies (Kolar & Lodge, 2001), and in particular to facilitate prevention and early detection, which are the most cost-effective for management (Simberloff *et al*., 2013). In addition, government and management agencies use lists of non-indigenous species in their policies to protect nature and reverse the degradation of ecosystems. For instance, the primary criterion for the descriptor D2 dedicated to non-indigenous species under the European “Marine Strategy Framework Directive” is the rate of novel introductions per 6-year period (European Commission *et al*., 2021). Comprehensive and accurate lists of non-indigenous species, their respective origin, and geographical and temporal spread are therefore crucial for an effective response and legislation to battle threats imposed by non-indigenous species. Unfortunately, the compilation of such lists is marred by the challenges involved (McGeoch *et al*., 2012; Costello *et al*., 2021). At local scales, lists may be confounded by limited occurrence data and hence underestimate the number and spread of non-indigenous species. At a more fundamental level, taxonomic uncertainty and the associated lack of expertise in species identification are regarded as severe problems (Zenetos *et al*., 2017). The effects of taxonomic uncertainty are likely more pronounced for less studied taxa and poorly sampled regions. For example, upon re-examination of about 100 potential non-indigenous taxa of marine molluscs, almost half of the records turned out to be misidentifications or the distributional data were incorrect (Zenetos *et al*., 2017). While DNA-assisted identification has the potential to solve identification problems, misidentifications of entries in genetic databases combined with geographic and taxonomic sampling bias make it a challenge in itself to correctly interpret gene sequence data (Viard *et al*., 2019; Fort *et al*., 2021; Tran *et al*., 2022). In addition, taxonomic knowledge is not static. Evolving taxonomic insights, often derived from genetic and biogeographic studies, alter our views on the indigenous or non-indigenous nature of taxa, requiring checklists to be continuously updated (Taylor, 2010; Guareschi & Wood, 2019). This problem is exacerbated in the marine environment where many cryptic species have been documented (Appeltans *et al*., 2012).

The above-mentioned problems related to lists of non-indigenous species definitely apply to seaweeds, which represent one of the largest groups of marine non-indigenous organisms, constituting between 20 and 29% of all marine non-indigenous species in the Northeast Atlantic Ocean, the Mediterranean Sea and Macaronesia (hereafter referred to as “the study area”) (Schaffelke *et al*., 2006; Molnar *et al*., 2008; Katsanevakis *et al*., 2013) (Fig. 1; Fig. 2). The consequences of non-indigenous species on native ecosystems have only been studied in a very limited number of species. Although some non-indigenous species have been observed to have positive ecosystem effects (e.g. *Gracilaria vermiculophylla* in the Venice Lagoon and Northeast Atlantic mudflats; Davoult *et al*., 2017; Sfriso, 2020), impact studies on such seaweeds have mostly detected negative ecological effects, with reduction in abundance of native biota being most frequently reported (Williams & Smith, 2007; Weinberger *et al*., 2008; Hammann *et al*., 2013; Katsanevakis *et al*., 2014; Maggi *et al*., 2015; Bulleri *et al*., 2017; Anton *et al*., 2019). However, contrary to the evidence of substantial negative impact on coastal ecosystems of many non-indigenous seaweeds (e.g. *Caulerpa cylindracea*, *Caulerpa taxifolia*, *Codium fragile*), so far *Rugulopteryx okamurae* is the only seaweed included in the list of invasive alien species of Union concern (COMMISSION IMPLEMENTING REGULATION (EU) 2022/1203 of 12 July 2022 amending Implementing Regulation (EU) 2016/1141). This EU regulation enforces member states to adopt measures to prevent, minimise or mitigate the adverse impact of those species.

**FIGURE 1.**
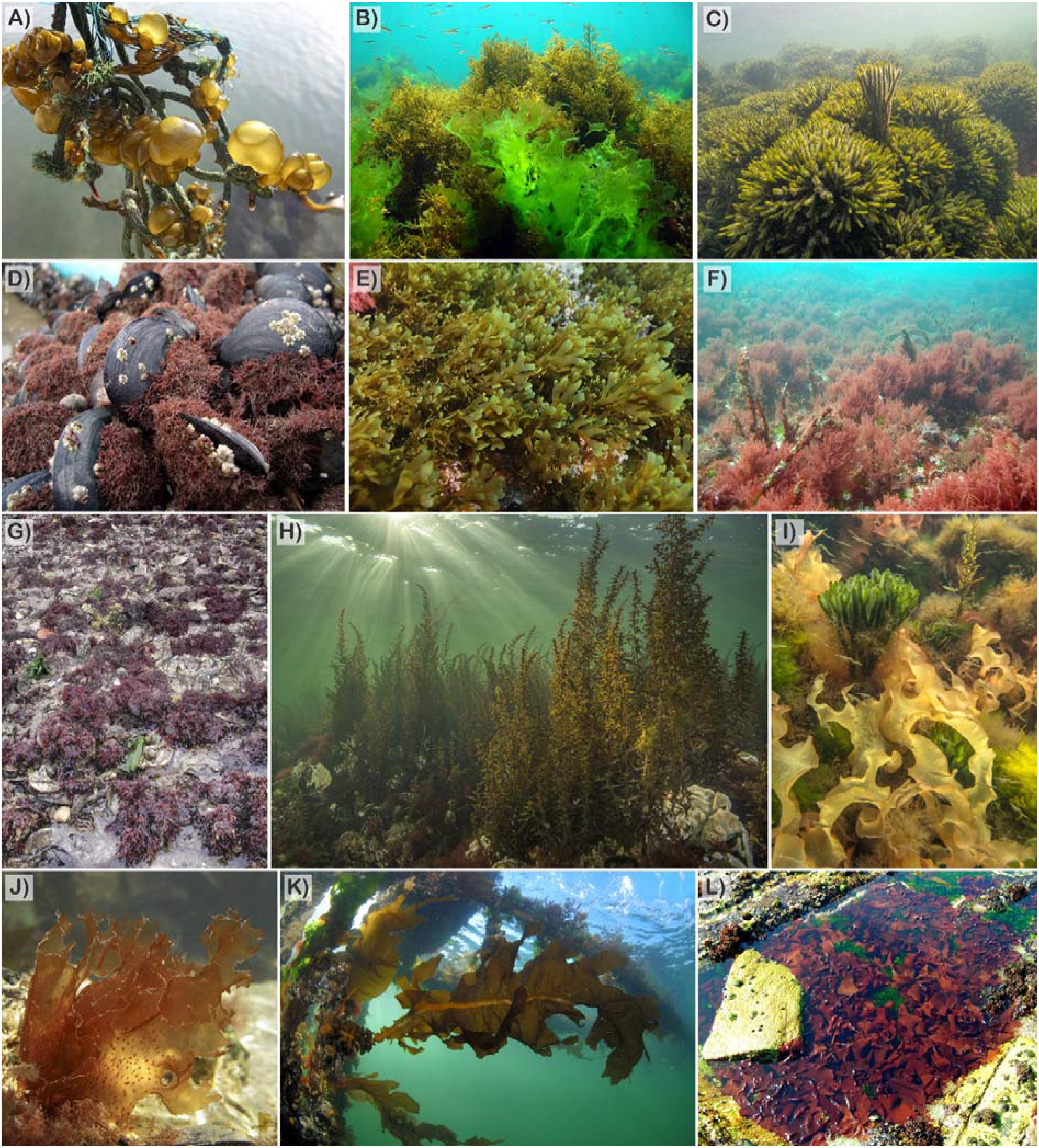
Illustration of selected non-indigenous species in the study area. A) *Colpomenia peregrina* growing attached to nylon fishing net in a harbour (photo: Frank Perk, the Netherlands); The green non-indigenous species *Ulva australis* (photo: Ignacio Bárbara, Atlantic coast Spain); A dense *Codium fragile* subsp. *fragile* reef (photo: Mick Otten, the Netherlands); D) *Caulacanthus okamurae* often grows high in the intertidal (photo: Ignacio Bárbara, Atlantic coast Spain); E) *Rugulopteryx okamurae* has been introduced in the Northeast Atlantic Ocean, Mediterranean Sea and in Macaronesia (photo: Sandrine Ruitton, Mediterranean coast France); F) *Asparagopsis armata* is often regarded as a high-nuisance invasive species (photo: Ignacio Bárbara, Atlantic coast Spain); G) *Gelidium vagum* has only been reported from the Netherlands but is locally very abundant (photo: Mart Karremans, the Netherlands); H) A *Sargassum muticum* forest (photo: Rob Aarsen, the Netherlands); I) *Grateloupia turuturu*, *Codium fragile* subsp. *fragile*, and *Sargassum muticum* covering the seabed (photo: Ad Aleman, the Netherlands); J) A fertile specimen of *Nitophyllum stellatocorticatum* (photo: Mart Karremans, the Netherlands); K) *Undaria pinnatifida* growing attached to aquaculture facilities (photo: Ron Offermans, the Netherlands); L) A tidal pool with *Grateloupia turuturu* (photo: Ignacio Bárbara, Atlantic coast of Spain).

**FIGURE 2.**
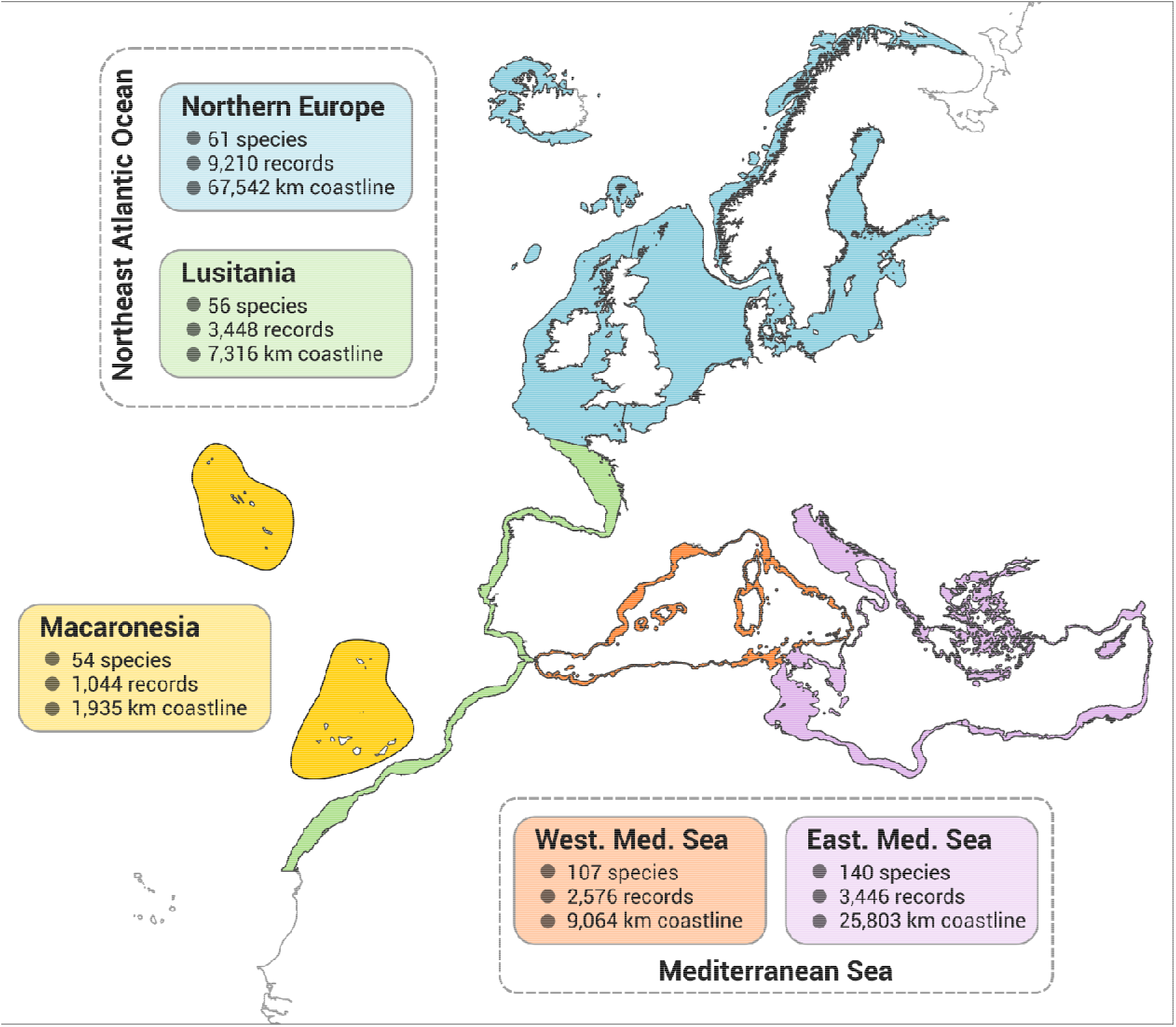
Flowchart for assessing the taxonomic confidence and biogeographic status of putative non-indigenous species. This figure builds on the concepts from Essl *et al*. (2018)

Regional lists of non-indigenous seaweed species have been regularly published until recently. For the Mediterranean Sea, which has been disproportionately affected by non-indigenous species as well as other stressors (Lejeusne *et al*., 2010; Katsanevakis *et al*., 2014), non-indigenous seaweeds have been critically revised on a regular basis (Verlaque *et al*., 2015; Zenetos *et al*., 2017; Galil *et al*., 2021). Non-indigenous seaweeds of Macaronesia were included in Borges *et al*. (2010), Chainho *et al*. (2015), Gallardo *et al*. (2016) and Castro *et al*. (2022). Bárbara *et al*. (2005) and Brodie *et al*. (2016) provided a list of non-indigenous seaweeds as part of a revised check-list of Galician and British seaweeds, respectively. However, there are gaps and uncertainties for some regions, and more importantly, a critical compilation encompassing the Northeast Atlantic Ocean, Macaronesian and Mediterranean regions is currently lacking. The absence of a critically revised list in the study area not only impedes a comprehensive overview of non-indigenous seaweeds, but may also introduce ambiguity related to the status of specific taxa due to differences in the criteria used to define non-indigenous species (see Materials and Methods). In addition, in the absence of a comprehensive list, spatial and temporal patterns of introductions are difficult to deduce.

To address this knowledge gap, we compiled a database of non-indigenous seaweeds in the Northeast Atlantic Ocean, the Mediterranean Sea and Macaronesia with their distribution records, their likely origin and putative introduction vectors. These data are used to provide a quantitative assessment of the spatio-temporal dynamics of primary and secondary introductions and to detect shortcomings in the monitoring and legislation required to tackle the introduction of non-indigenous species more effectively.

## Materials and methods

### Data compilation

We compiled a database of non-indigenous marine seaweed species records from three regions, namely the Northeast Atlantic Ocean (excluding Greenland), the Mediterranean Sea and Macaronesia (Fig. 2). For some of the analyses we subdivided the Northeast Atlantic Ocean into Lusitania and Northern Europe and the Mediterranean Sea into a Western and Eastern part. With respect to Macaronesia, the compilation includes records from the Azores, Canary Islands, Madeira and the Salvagen Islands, but not Cape Verde. The dataset builds on previous lists by Mineur *et al*. (2010) and Verlaque *et al*. (2015), and includes published records of species occurring in a natural environment and flagged as non-indigenous in the study area irrespective of taxonomic confidence and biogeographic status (see below). In addition, we included unpublished records produced by various research projects conducted by, amongst others, the Station Biologique de Roscoff (France), National Biodiversity Data Centre (Ireland), Stichting ANEMOON (the Netherlands), Scottish Natural Heritage (Scotland), the ICES Working Group on Introductions and Transfers of Marine Organisms 2004, as well as the European Alien Species Information Network (EASIN, 2022) records, collection data, GBIF records and personal data. All records were added to the database under the name they were reported as. Names were updated according to the most recent taxonomic consensus (AlgaeBase, Guiry & Guiry, 2023).

The species listed as non-indigenous include those that are naturalised (i.e. having established permanent, self-maintaining populations), as well as species for which no information is available on population status (i.e. species referred to as ‘alien’ by Verlaque *et al*. 2015). Species that have been demonstrated to be misidentifications or unsupported records are excluded from the list. To promote consistency in definitions and criteria used to determine whether a species is non-indigenous, we have adopted the criteria for assessing the biogeographic status proposed by Essl *et al*. (2018) (Fig. 3). This framework stresses 1) the need for crossing a biogeographic barrier, 2) the involvement of direct or indirect human agencies in the physical movement of individuals, spores or fragments, and 3) the ability of the species to reproduce without human assistance in the introduced range. The combination of these criteria excludes records of species which are in the process of expanding their range naturally, for example as a result of global warming. Species entering the Mediterranean Sea through the Suez Canal (i.e. Lessepsian migrants), on the other hand are considered non-indigenous because of the anthropogenic nature of the dispersal corridor. In contrary, species entering the Mediterranean Sea through the strait of Gibraltar, without a human-vector, are not considered as non-indigenous. The dataset also includes species indigenous to the study area that have demonstrably become displaced within he study area as a result of human-mediated exchanges. Examples include exchanges of species between Atlantic and Mediterranean shores. Species for which the area of origin is unknown are assigned as ‘cryptogenic’ (sensu Carlton, 1996). In cases where there is not sufficient information to be conclusive on their biogeographic status, species are labelled as ‘data deficient’. Species with low uncertainty, for which there is no doubt about their non-indigenous status, have been labelled ‘non-indigenous’.

**FIGURE 3.**
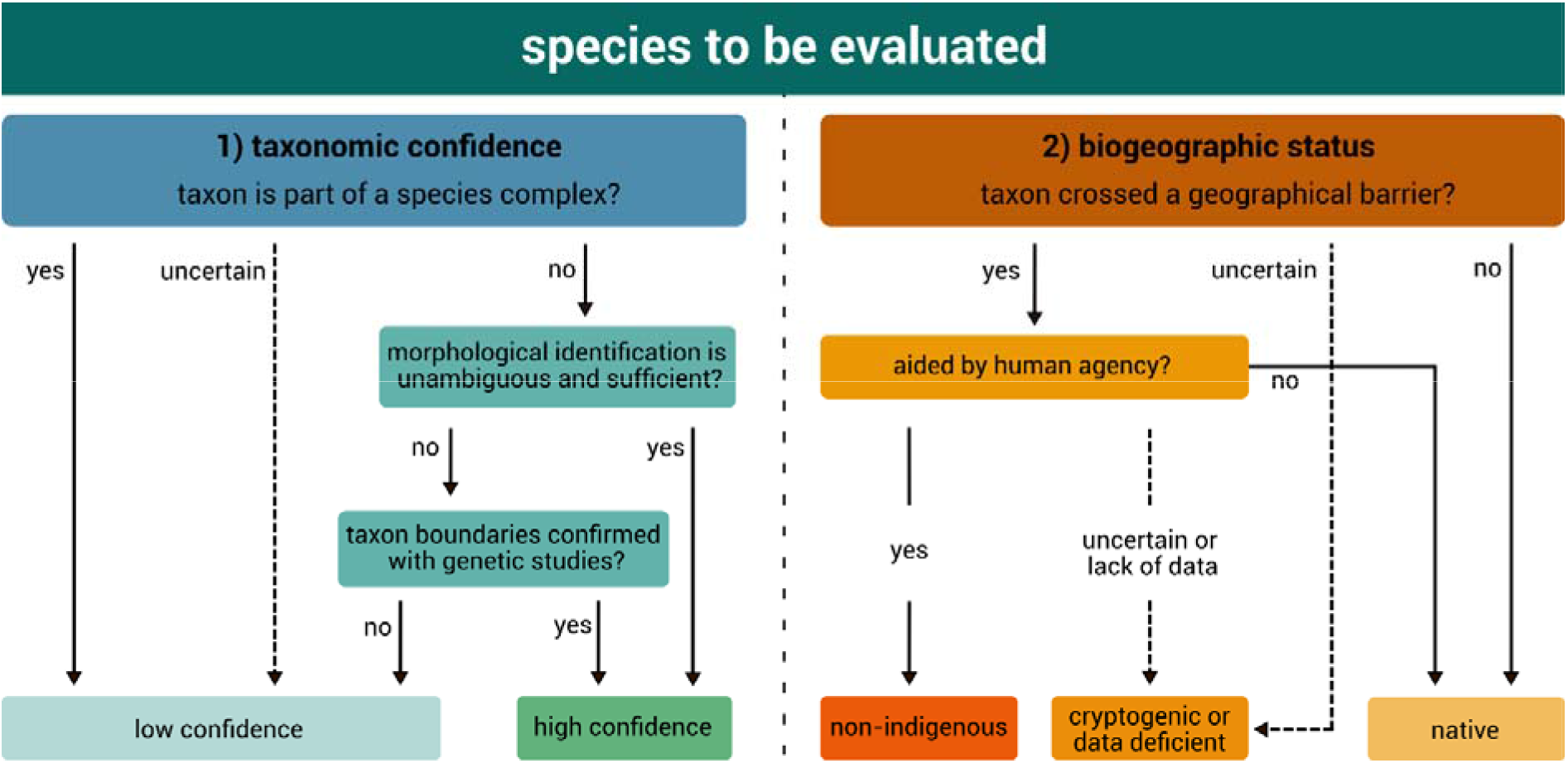
Map of the study area (Northern Europe, Lusitania, Macaronesia, Western Mediterranean Sea, and Eastern Mediterranean Sea), with indication of the number of recorded non-indigenous species, number of records and the length of the coastline. The number of species includes cryptogenic and data deficient species.

Added to these criteria but highly relevant with respect to seaweeds, where a solid taxonomic framework is often lacking for many taxa, is taxonomic confidence. We assigned a ‘high-confidence score’ (score = 1) to accepted nominal species that had not been shown to be a species complex based on molecular studies in their introduced or native ranges. A high score was also assigned to species for which, so far, there is sufficient confidence in unambiguous identification based on morphology. We acknowledge, however, that the latter does not rule out the potential existence of cryptic species hidden under the accepted nominal species. Conversely, species that belong to an understudied complex of cryptic species were assigned a low-confidence score (score = 0). A cryptic species is defined here as a taxon composed of two or more species that have been classified as a single nominal species, because they were initially not distinguished based on their morphological characteristics (Bickford *et al*., 2007; Pante *et al*., 2015).

Recognising we cannot be conclusive about the non-indigenous status of many seaweed species, we explicitly acknowledge the uncertainty in the assessment of the taxonomic as well as biogeographic status of putative non-indigenous seaweeds in the study area (Fig. 3). The status of each species is concisely described in Suppl. Material Table S1.

For every species we determined the year when the species was first reported in the Northeast Atlantic Ocean, Mediterranean Sea and/or Macaronesia. Where possible, this date refers to the year the species was detected (i.e. collection date) rather than when the record was published (i.e. publication date). We acknowledge that detection dates may not portray the actual date the species was introduced. For each species an estimate is provided for its native biogeographic range. If the native range could not be assessed, we indicated ‘uncertain’. The putative distribution of the species was based on literature reports included in AlgaeBase (Guiry & Guiry, 2023). Species traits (e.g. thallus size) were obtained from AlgaeTraits (Vranken *et al*., 2022). For spatial and temporal analyses, distribution records were filtered on a combination of unique year, coordinates and species name to eliminate potential duplicate records. The complete dataset has been archived at Zenodo and is available at DOI: 10.5281/zenodo.7798640. This dataset contains the following information for each record: currently accepted scientific name, the scientific name under which it was originally reported, year of record, location, country, coordinates and reference.

## Results and Discussion

A total of 19,724 records of non-indigenous seaweeds were collected dating from 1808 to 2022 (Fig. 2). Of these, 17,104 were retained after removing duplicates and incomplete data. The geographical distribution of the records highlights considerable sampling of non-indigenous seaweeds from all coastlines in the study area (Fig. 2). The list contains 227 species (Table 1). Non-indigenous species make up approximately 10% of the seaweed flora in the Mediterranean Sea, 6% in the Northeast Atlantic Ocean and 4% of the Macaronesian flora. The total number of 227 includes all species regardless of taxonomic and biogeographic uncertainty. For 84 species neither their non-indigenous status nor their taxonomy is challenged (Fig. 4). These species make up 83% of the distribution records in the database. Half of the distribution records can be attributed to only five species (*Sargassum muticum*, *Bonnemaisonia hamifera*, *Asparagopsis armata*, *Caulerpa cylindracea* and *Colpomenia peregrina*). Fifty-six species are most likely non-indigenous, but decisions are hampered by taxonomic uncertainties (Fig. 4). On the other hand, 87 of the 227 species have a cryptogenic or data deficient status (30 species with an uncertain biogeographic status, and 57 species for which both the geographic status and taxonomic confidence are uncertain), meaning that the evidence for a non-indigenous status is mediocre to weak (Fig. 4).

**FIGURE 4.**
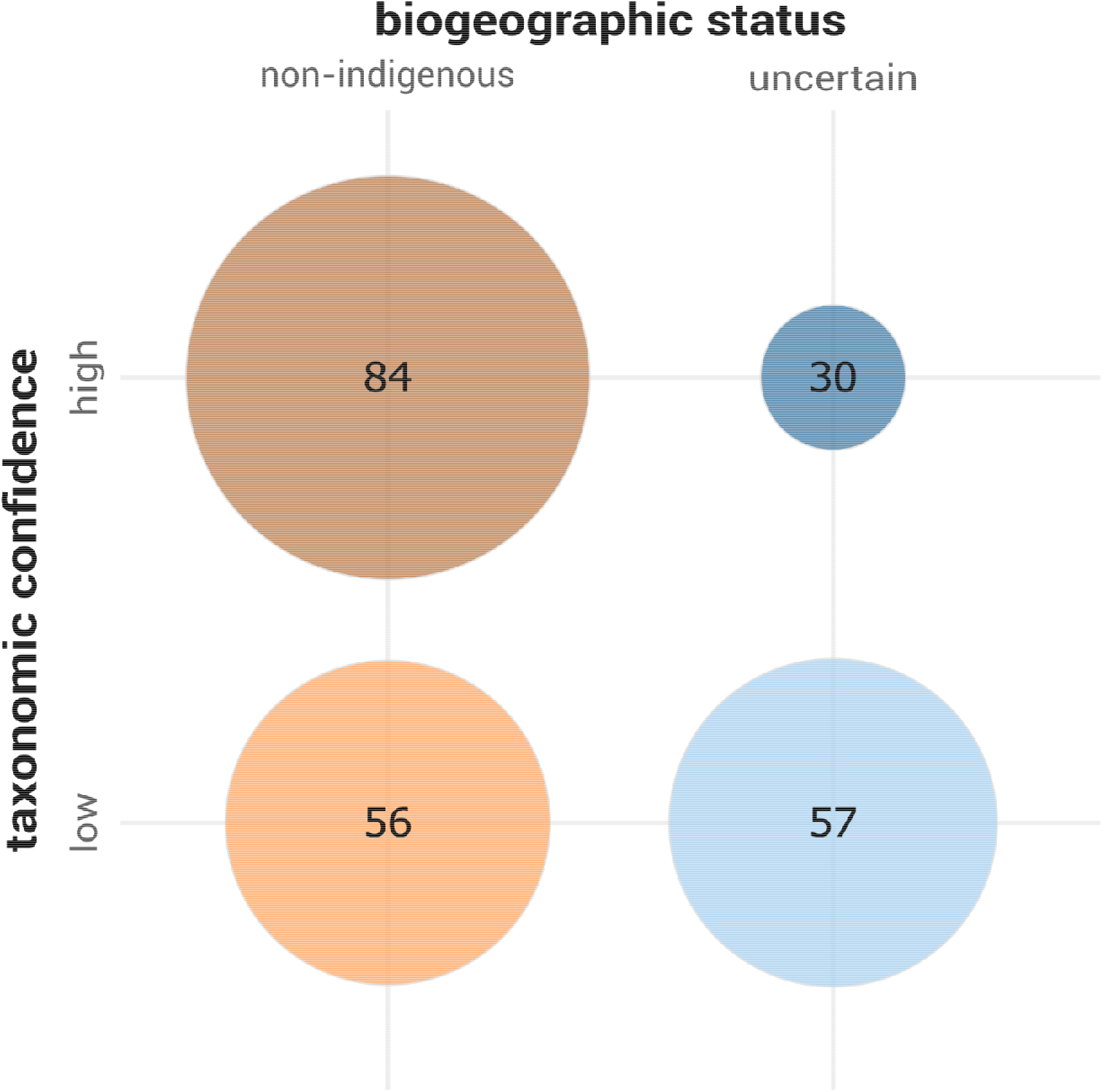
Biogeographic status (either non-indigenous or uncertain, the latter category including the ‘cryptogenic’ and ‘data deficient’ species) and taxonomic confidence of seaweed species flagged as non-indigenous in the study area. Circle surface area corresponds to the number of species.

**Table 1.**
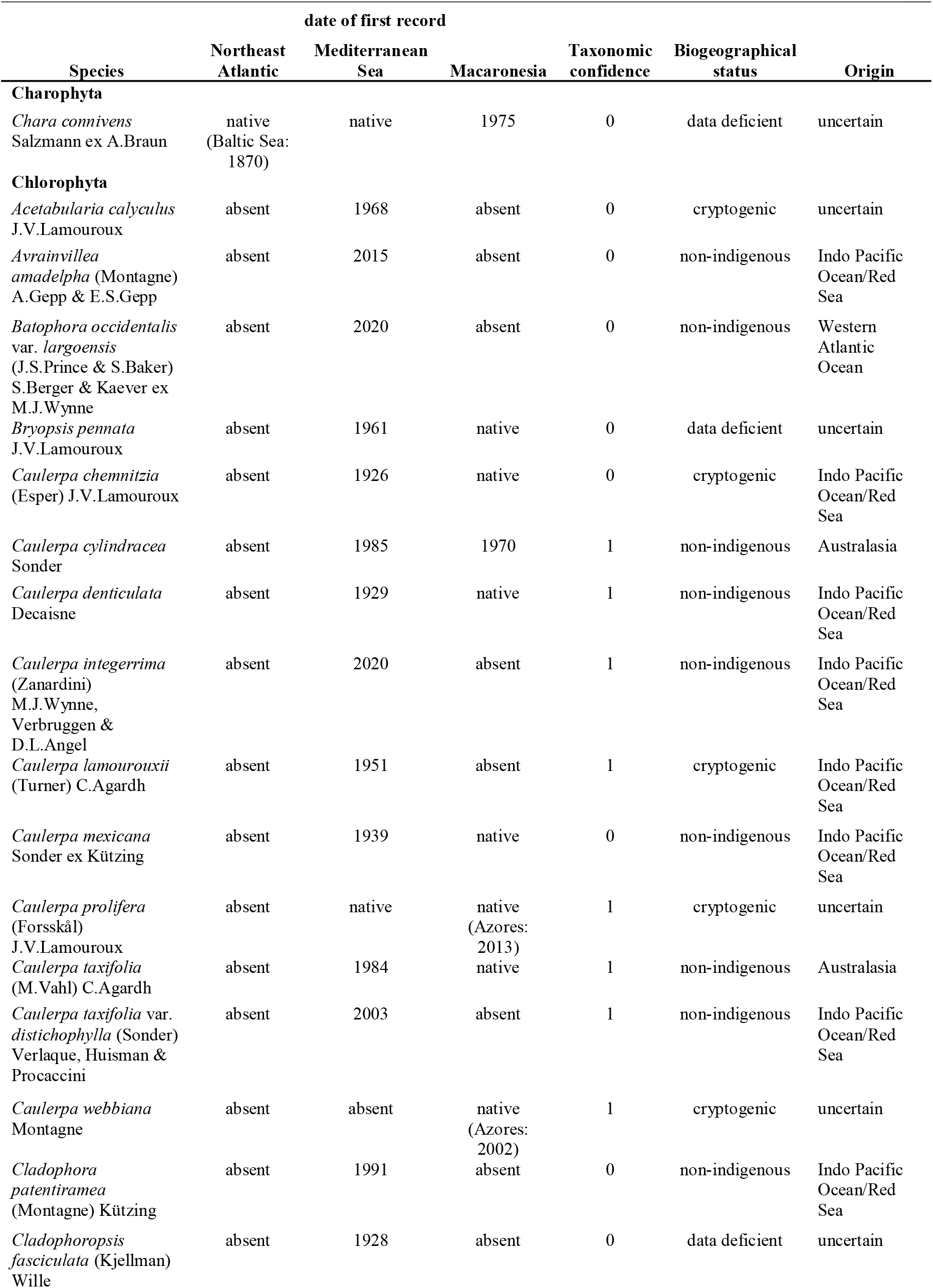
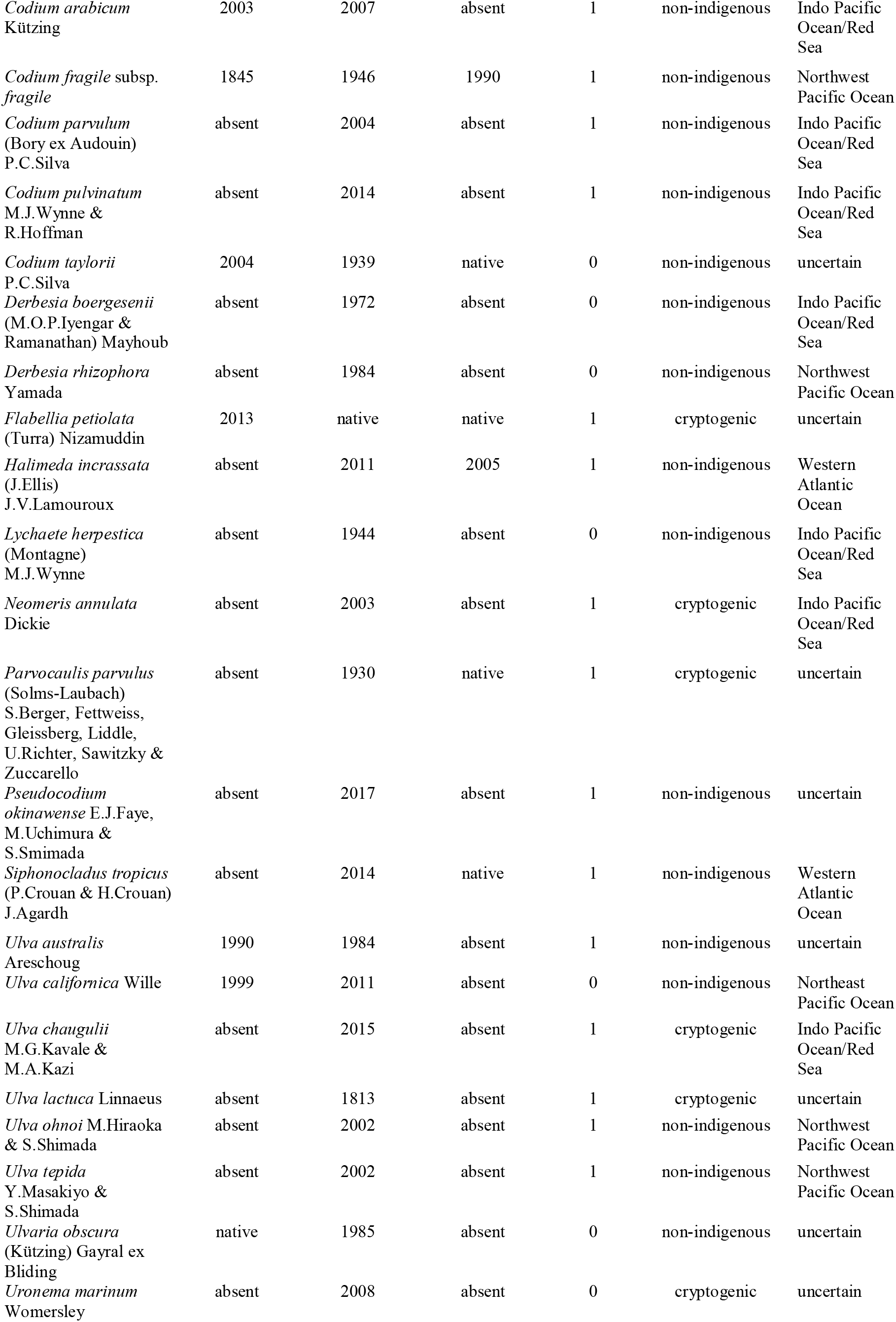
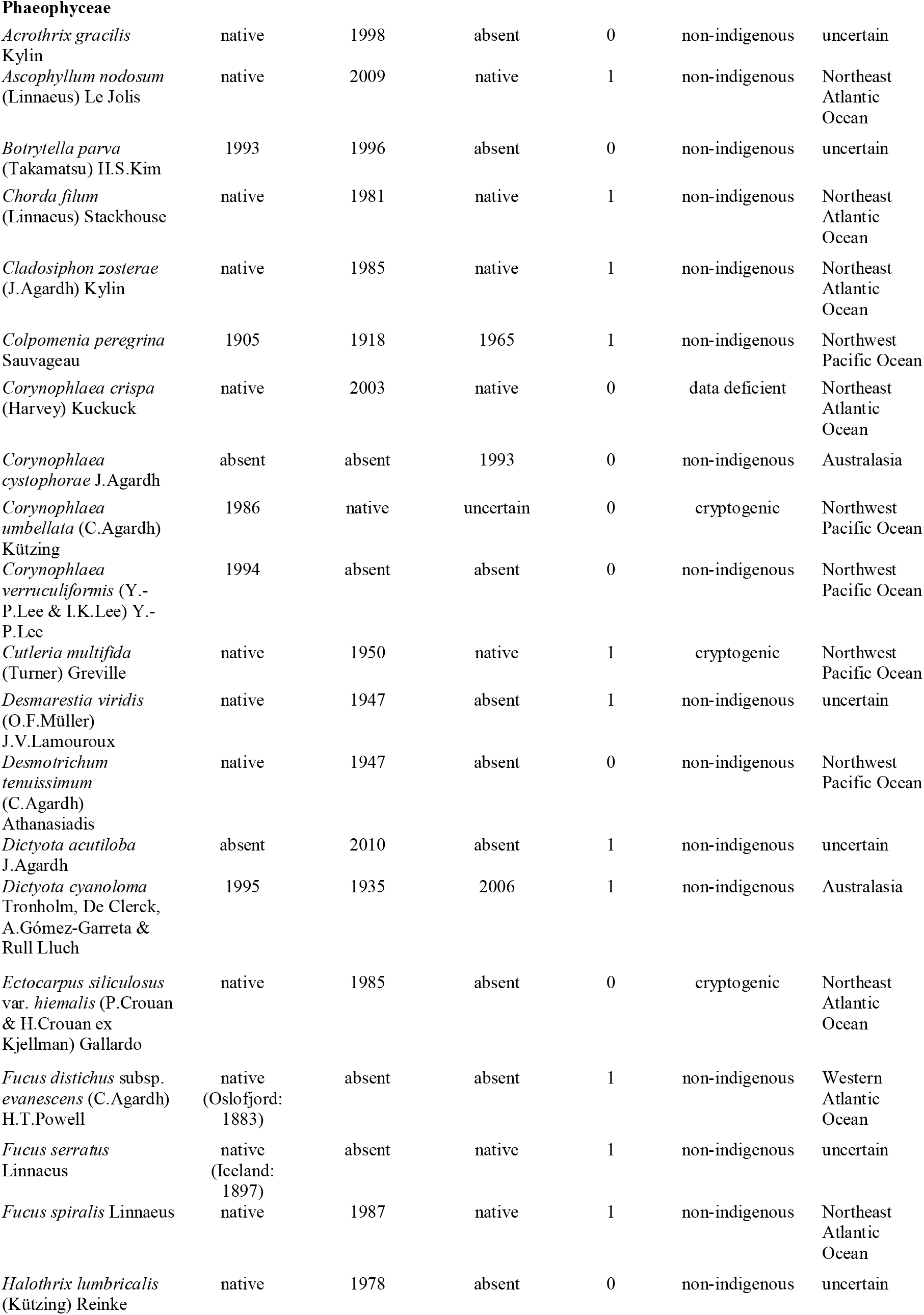
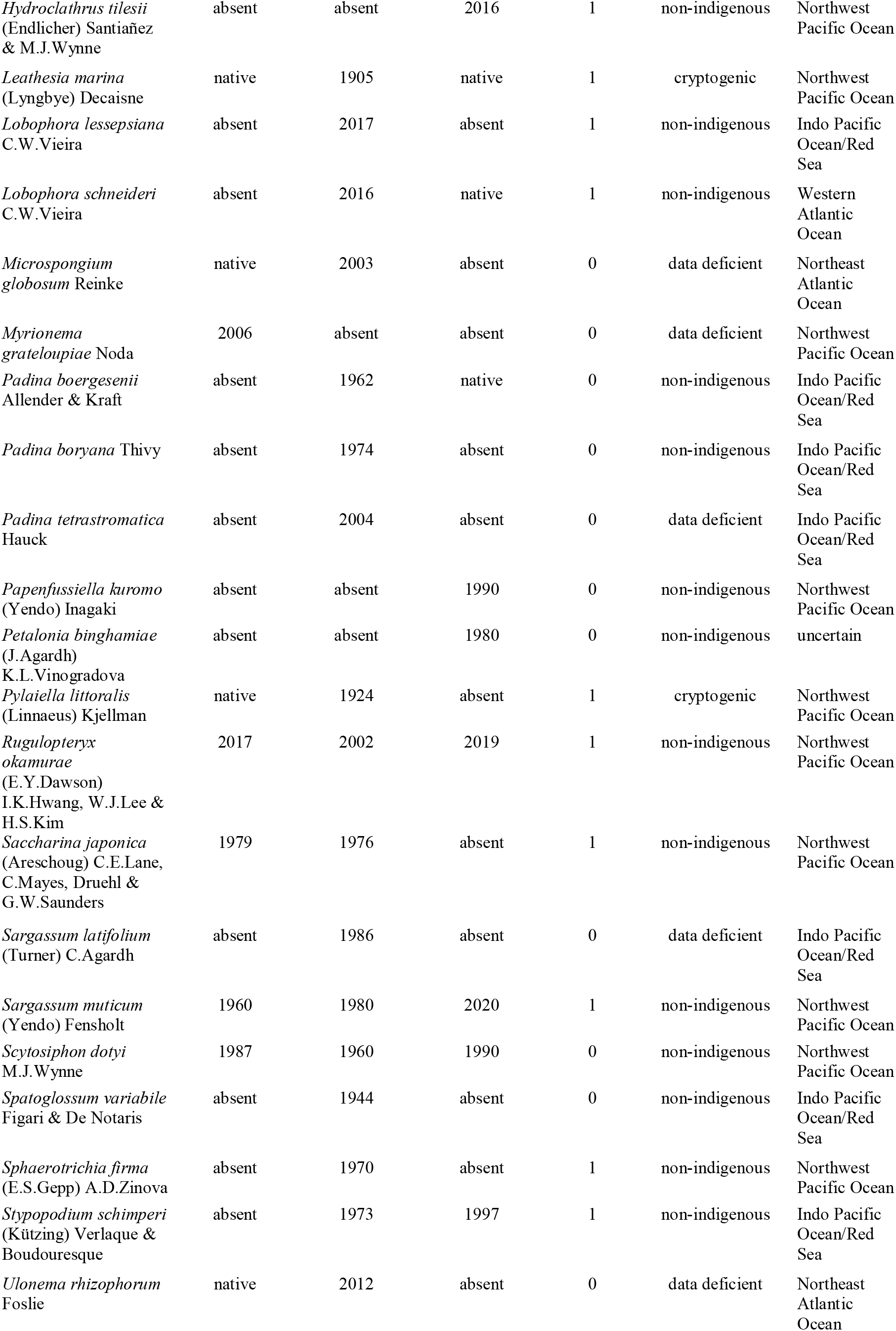
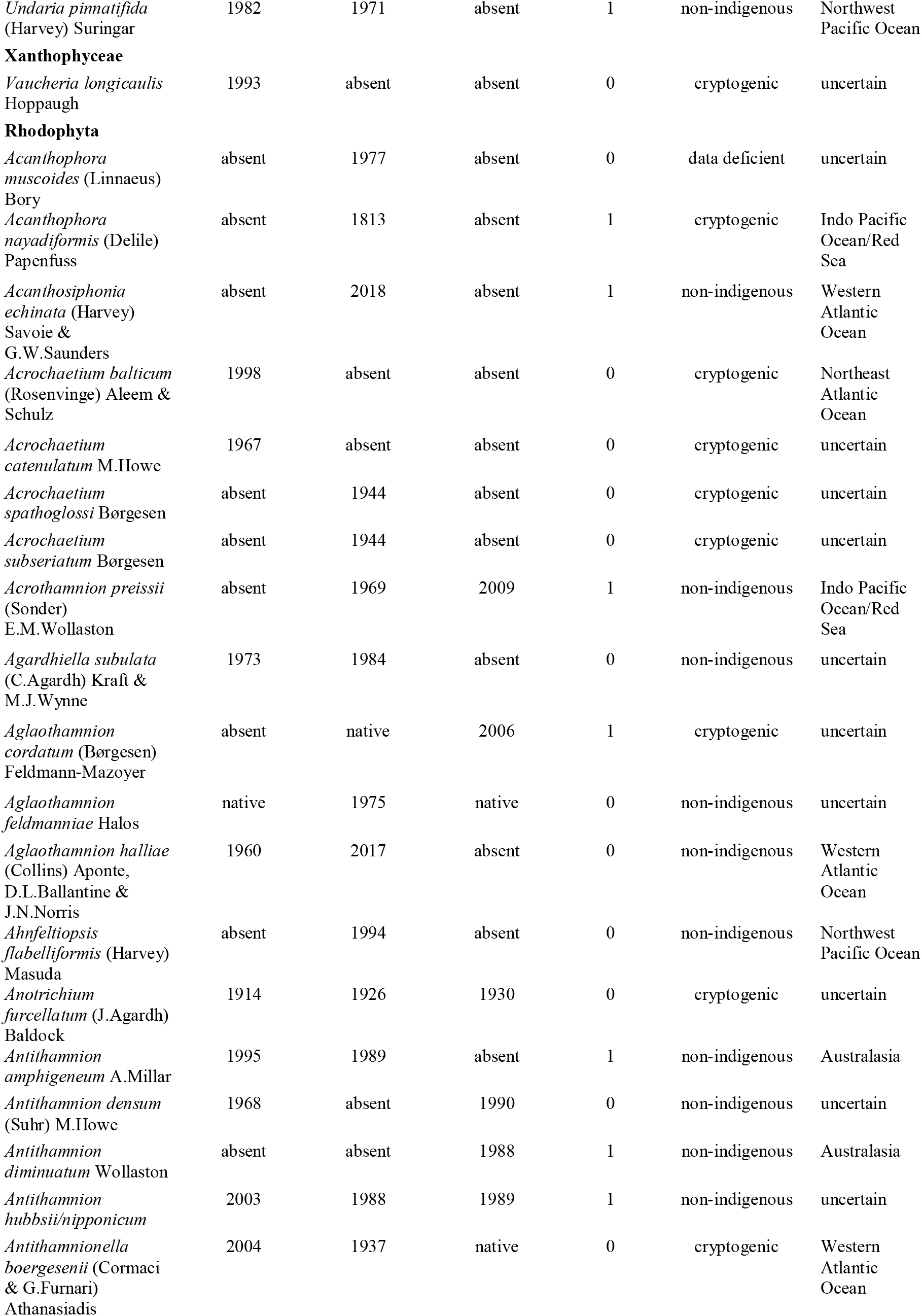
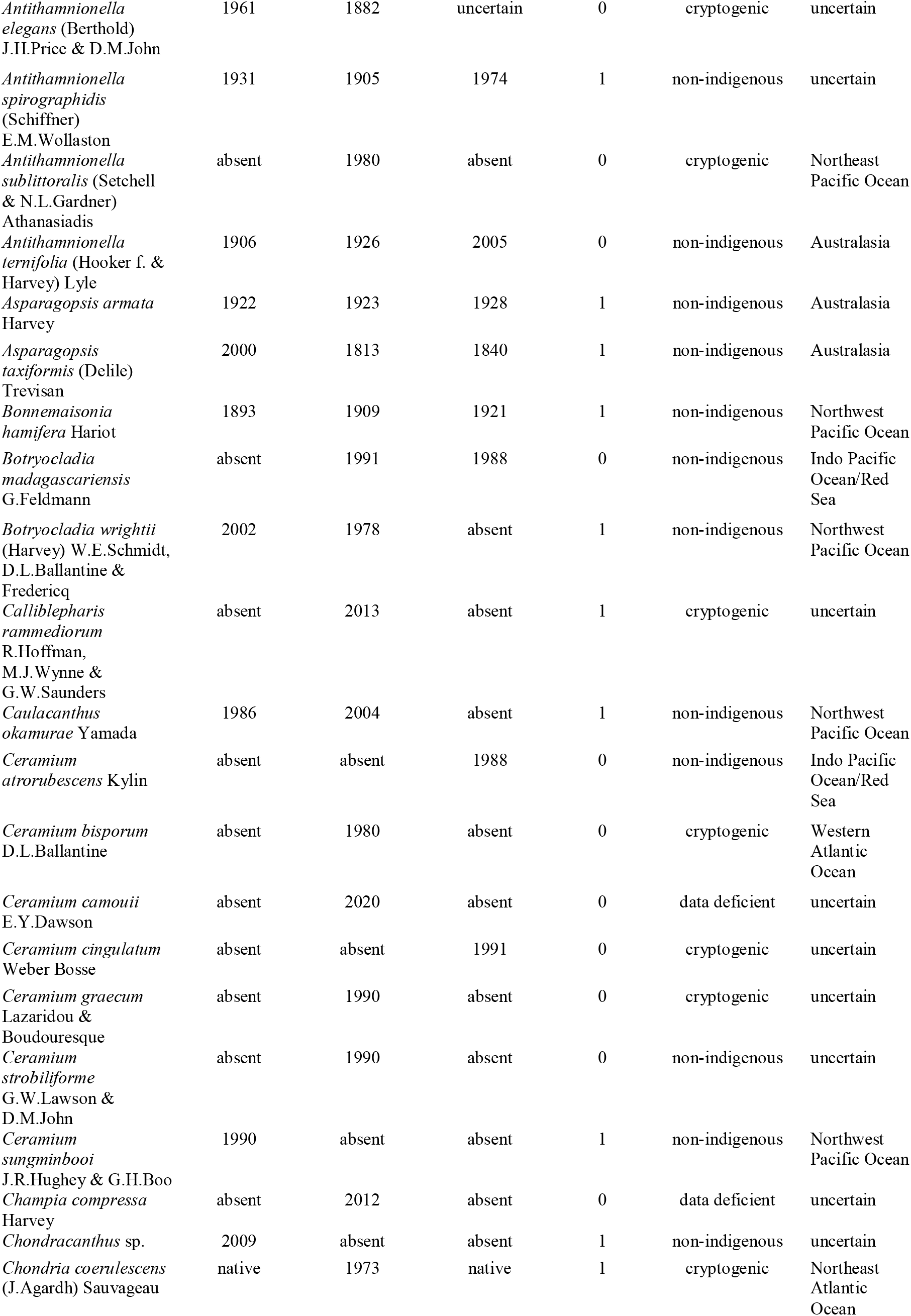
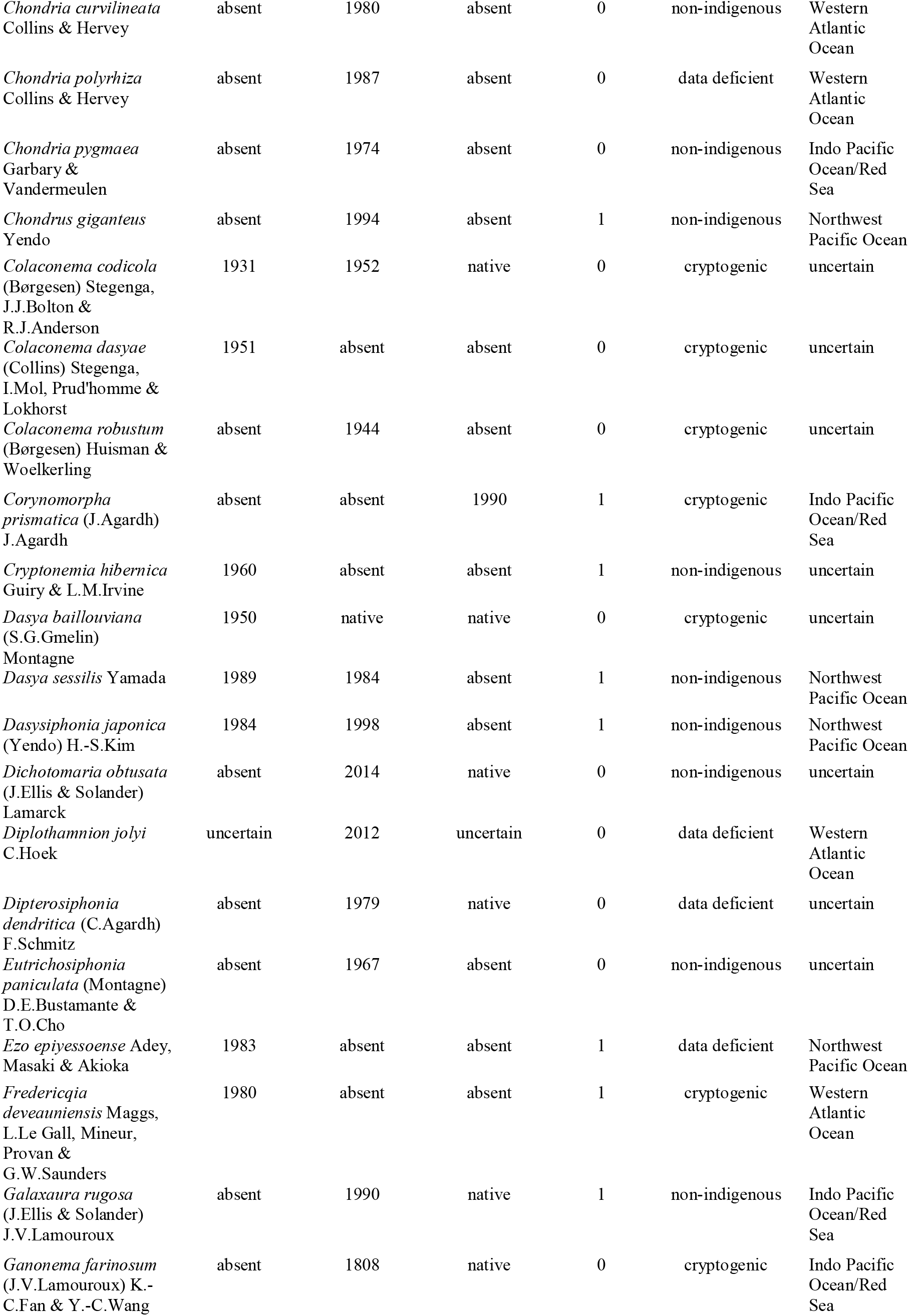
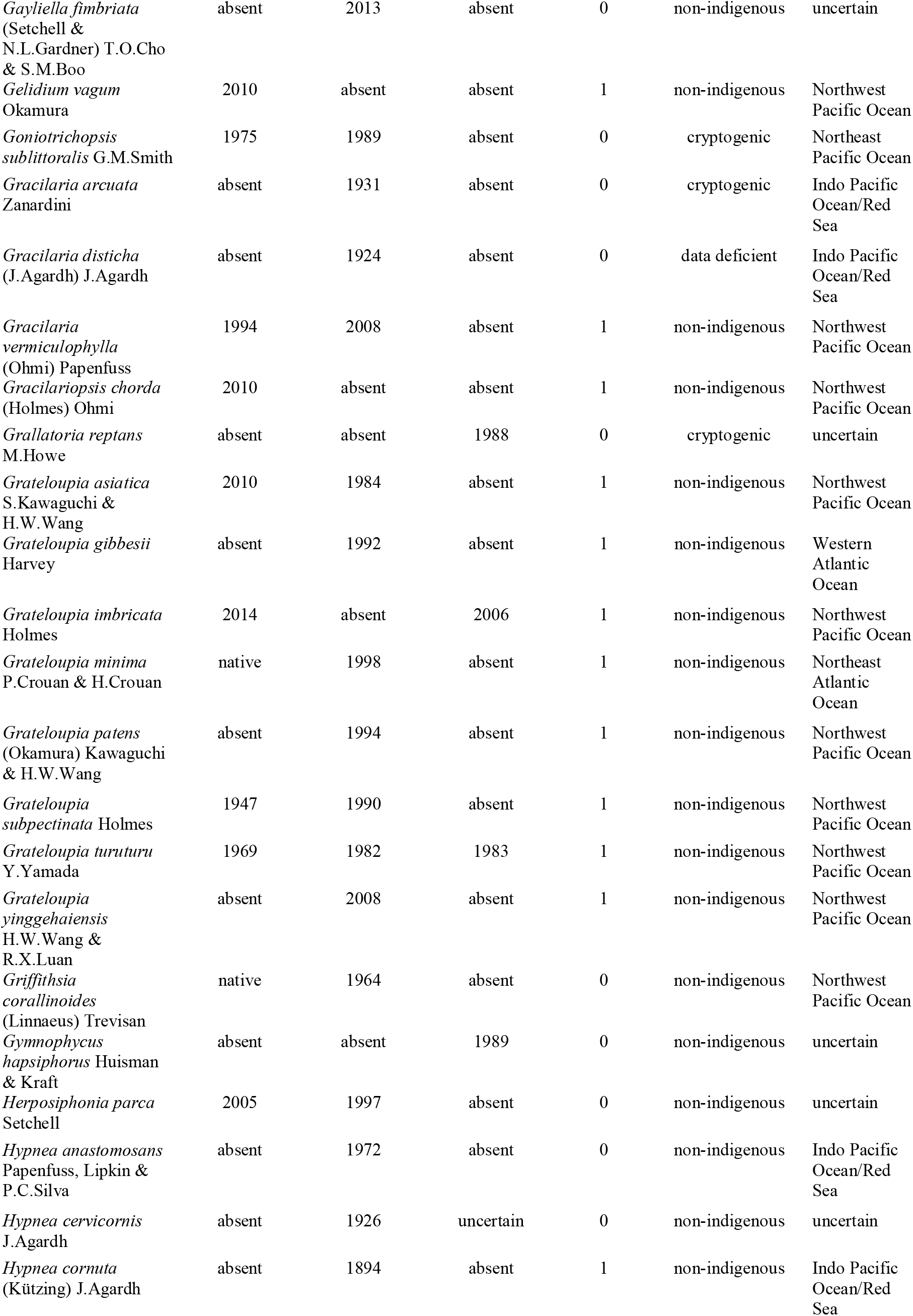
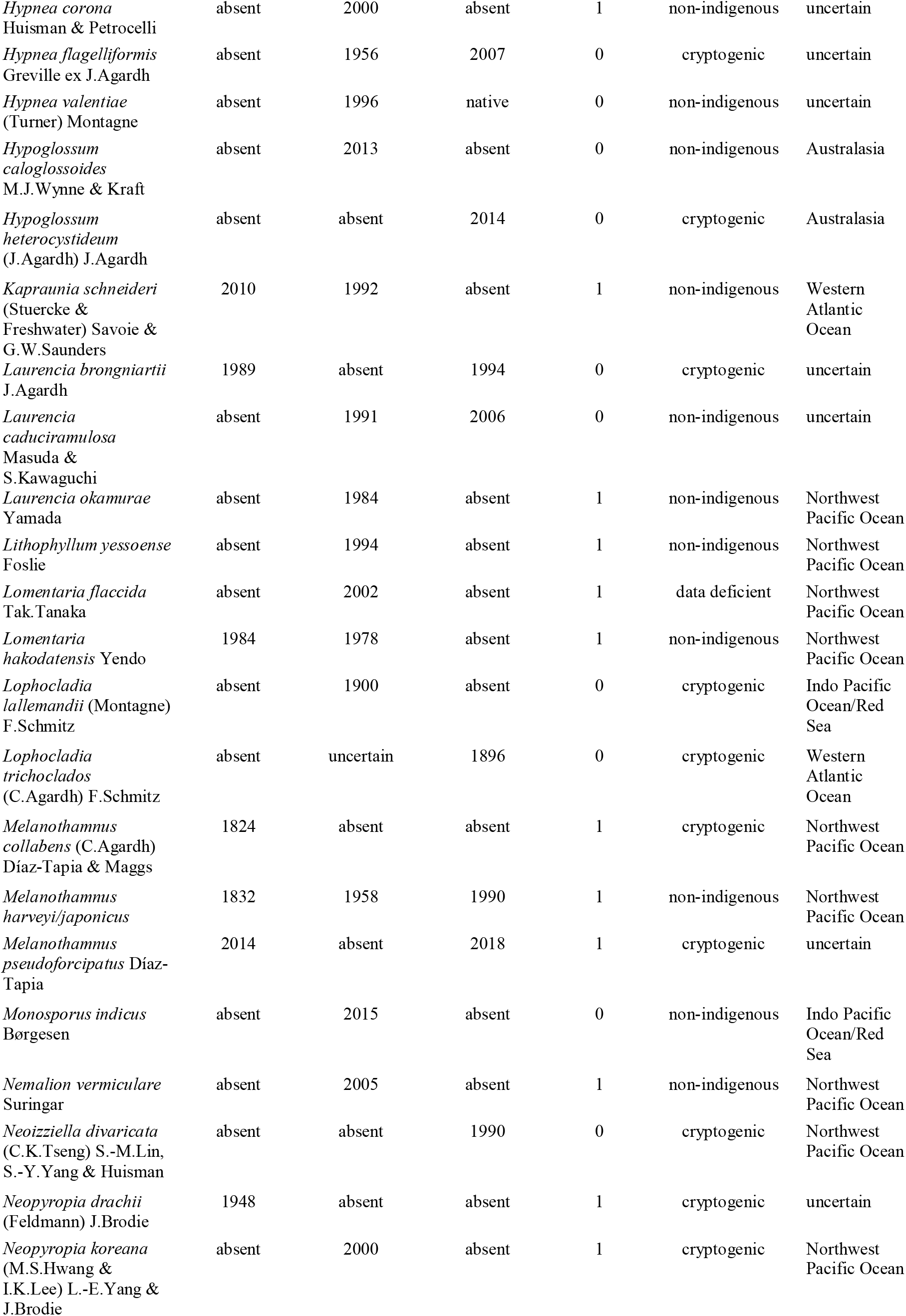
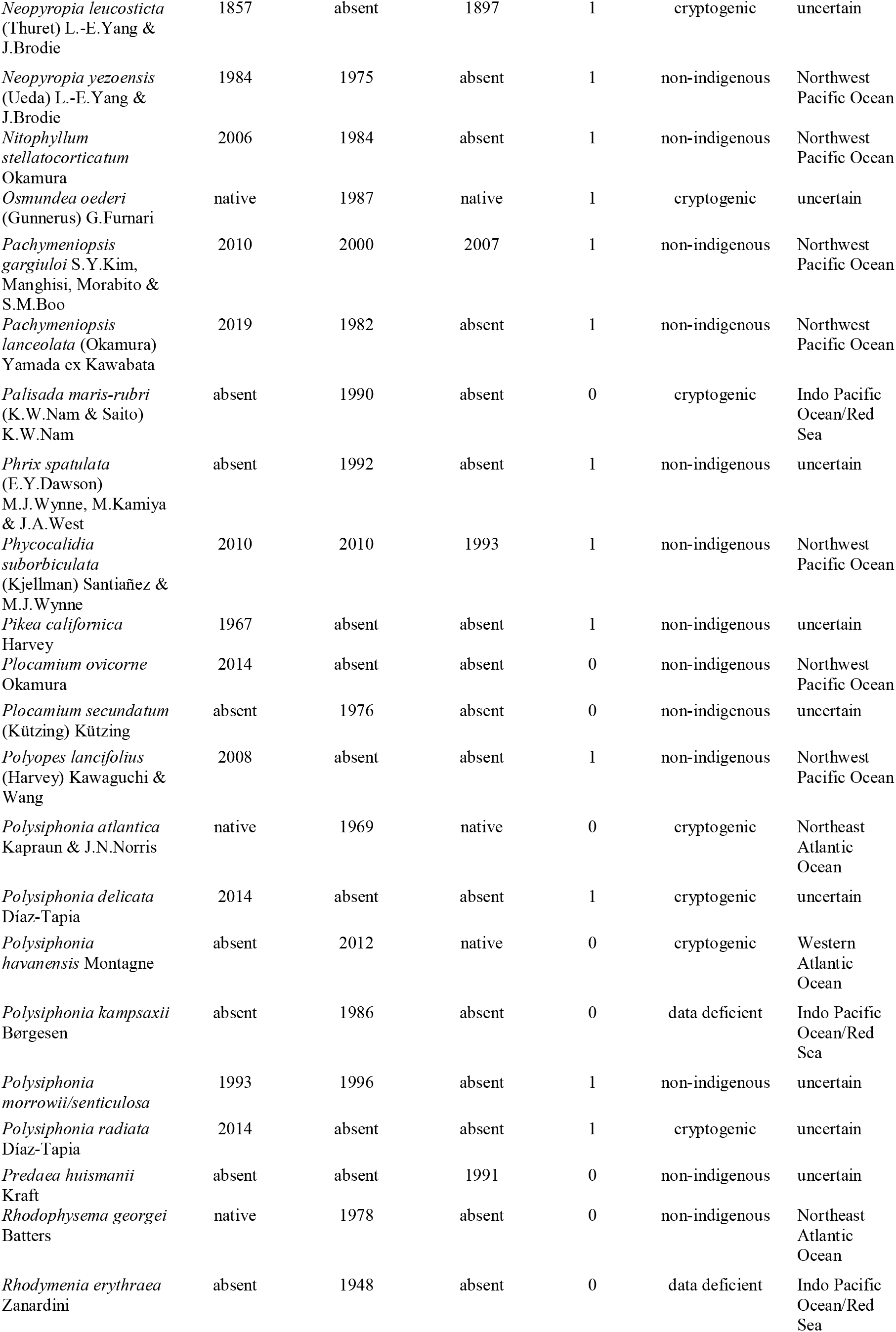
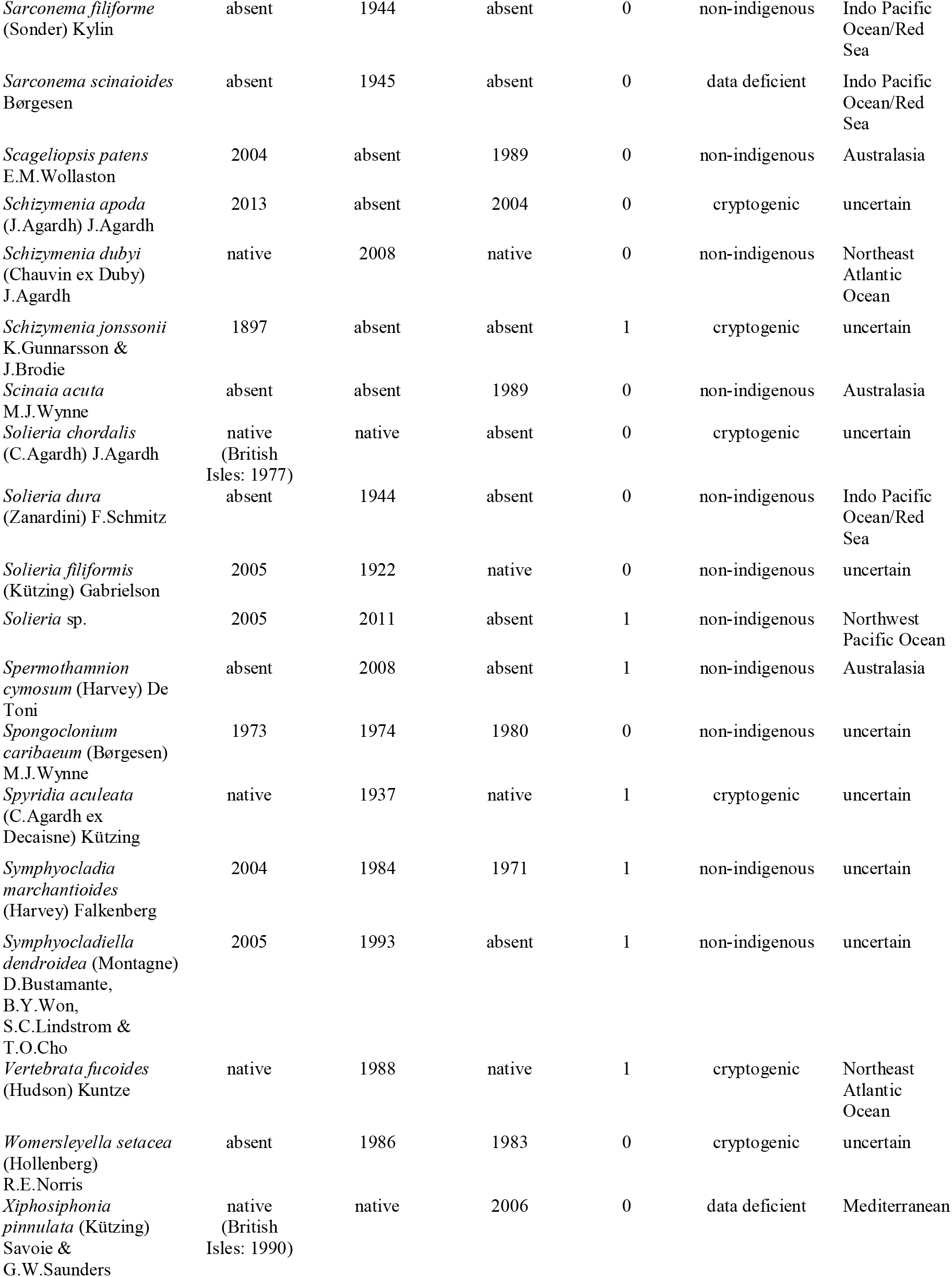
Overview of the non-indigenous seaweed species (Charophyta, Chlorophyta, Phaeophyceae, Xanthophyceae, Rhodophyta) reported from the study area.

We emphasise that the categorisation of putative non-indigenous species according to biogeographic uncertainty and taxonomic confidence emerged as a consensus among the authors of this paper. A literature search will undoubtedly reveal several additional species names that could potentially be added to the list of cryptogenic or data deficient species. However, there is little added value in incorporating species names which are wholly unsupported or most likely result from misidentifications or other mistakes. Evidently, both taxonomic and biogeographic uncertainty plague the compilation of lists and databases of non-indigenous species. Below we discuss how the level of sophistication of systematic and biogeographic knowledge translates to uncertainty in the number of non-indigenous seaweeds in the study area.

## Taxonomic confidence

For 140 species the non-indigenous nature of the species itself is not disputed. However, the reliable identification of 56 of those species is challenging, and therefore their current distribution as well as their putative region of origin are questionable. In most cases this uncertainty can be attributed to a poorly established taxonomic framework. Taxonomic uncertainty is rife in seaweeds. In the absence of DNA sequence data the identification of many seaweed species is particularly difficult (e.g. Van Oppen *et al*., 1996; Maggs *et al*., 2007; Cianciola *et al*., 2010; De Clerck *et al*., 2013; Verbruggen, 2014). Taxonomic uncertainty is much higher among small-sized species: 67% of species smaller than 5 cm are flagged as taxonomically uncertain, compared to 34% of species larger than 5 cm. Of the taxa larger than 5 cm with high taxonomic uncertainty are many that belong to genera that are widespread in tropical and warm-temperate regions (e.g. *Avrainvillea, Caulerpa, Codium, Dichotomaria, Ganonema, Hypnea*). From a biogeographic perspective, taxonomic uncertainty plagues “only” 20% of species with a Northwest Pacific origin (11 of 56 species), but 62% of species with a likely Lessepsian or tropical Indo-Pacific origin (26 of 42 species) (Table 1).

Recent advances in the taxonomy of several genera, nearly always assisted by DNA sequence data, have demonstrated that many so-called wide-ranging (or cosmopolitan) seaweeds actually consist of species complexes of morphologically almost indistinguishable species (pseudocryptic species), or even truly cryptic species which are indistinguishable based on morphological criteria. The individual species are often confined to specific geographic areas (e.g. Won *et al*., 2009; Vieira *et al*., 2017; Diaz-Tapia *et al*., 2018; Leliaert *et al*., 2018; Diaz-Tapia *et al*., 2020). A more refined taxonomic framework therefore alters our understanding of the biogeography of the species in many cases and consequently our interpretation of their native versus non-indigenous status. The *Caulerpa racemosa* complex is highly representative of how evolving insights into species diversity alter our views of the taxa being non-indigenous in the study area. While initially *Caulerpa* specimens with vesiculate branchlets collected in the Mediterranean Sea were identified as *C. racemosa*, the latter proved to be a complex of at least eight species, three of which (*C. chemnitzia*, *C. cylindracea* and *C. requienii*) are currently considered non-indigenous in the Mediterranean Sea (Verlaque *et al*., 2000; Verlaque *et al*., 2003; Draisma *et al*., 2014; Verlaque *et al*., 2015). Similarly, a better understanding of the taxonomy of foliose *Grateloupia* species resulted not only in the recognition that *G. turuturu* was introduced in the study area from the Northwest Pacific, as opposed to *G. doryphora* whose distribution is likely to be restricted to the Pacific coast of South America (Gavio & Fredericq, 2002), but also revealed that so-called non-indigenous foliose *Grateloupia* species in the study area were actually a mixture of two non-indigenous species, *G. lanceolata* and *G. turuturu,* and a native species, *G. lanceola*, which had been regarded a synonym of *G. doryphora* (Verlaque *et al*., 2005; Figueroa *et al*., 2007).

In many other instances, however, conspecificity of populations from the native and non-native regions remains to be demonstrated. There are also examples where several non-indigenous species are thought to be conspecific by some authors but regarded as distinct species by others. For example, some authors consider *Antithamnion hubbsii* distinct from *A. nipponicum* (Athanasiadis, 1996), while others treat the former as a synonym of the latter (e.g. Kim & Lee, 2012). Similarly, records of *Polysiphonia morrowii* and *P. senticulosa* likely belong to the same species, even though both species are regarded as distinct (D’Archino *et al*., 2013; Stegenga & Karremans, 2015; Piñeiro-Corbeira *et al*., 2020). Given the widespread nature of cryptic and pseudocryptic diversity in seaweeds, continuous efforts of DNA-assisted identifications through Sanger sequencing will probably continue to revise our view on non-indigenous species.

Although DNA sequence data are in many cases a great help in verifying species identities, this does not mean DNA solves every single problem like a magic wand. Apart from reference sequences in repositories not being available or reliable, patterns of genetic divergence can be complicated and prone to different interpretations. For example, differences in the interpretation of genetic patterns and species boundaries in the genus *Melanothamnus* led to the recognition of a single species, *M. harveyi* s.l. (McIvor *et al*., 2001) or by contrast to the recognition of at least three separate species, including *M. akkeshiensis*, *M. japonicus* and *M. harveyi* s.s. (Savoie & Saunders, 2015). The narrower species concept would result in an interpretation whereby *M. harveyi* is native to the Northeast Atlantic Ocean rather than a non-indigenous species introduced to the study area from the Northwest Pacific Ocean. Under the alternative scenario which recognises a single genetically diverse species, *M. harveyi* is widely distributed globally with both cryptogenic and non-indigenous haplotypes in the study area (Piñeiro-Corbeira *et al*., 2019). One should note that despite the availability of a good number of sequences of these species/haplotypes, the potential native area of the species (Northwest Pacific Ocean) has been scarcely sampled. Therefore, it is still possible that *M. harveyi* s.s. can be present in this region but remained undetected. Distribution records of *M. harveyi* and *M. japonicus* are included as *M. harveyi/japonicus* in our dataset.

## Biogeographic uncertainty

A lack of baseline data with respect to the global distribution of seaweeds is the major contributor to biogeographic uncertainty reported for 87 taxa (Fig. 4). Brown and green seaweeds display slightly less biogeographic uncertainty, 28% and 34%, respectively, compared to 42% for red seaweeds. Baseline data of seaweed diversity along coastlines in the study area as well as the putative native regions in the form of herbarium collections, censuses and historical checklists can serve as a reference for the presence of species in a given area. Here as well, low confidence in the taxonomy and identification of seaweeds makes the interpretation of species lists exceedingly difficult. If a species is not reliably identified, its distribution is not reliable. As a result, biogeographic and taxonomic uncertainties usually go hand-in-hand. Three different categories of factors that lead to biogeographic uncertainty are discussed below.

### Pseudo-indigenous species

Several seaweed species have been described from the study area that were presumed native, but later turned out to be non-indigenous species. Carlton (2009) named such species pseudo-indigenous. For example, *Dictyota cyanoloma* was described as a new species from the Mediterranean Sea and Macaronesia (Tronholm *et al*., 2010), but subsequent collecting efforts revealed that the species most likely represents a cryptic introduction (Aragay Soler *et al*., 2016; Steen *et al*., 2017; Tran *et al*., 2021). Similarly, *Porphyra olivii* described from Greece (Brodie *et al*., 2007a) turned out to be conspecific with *Neopyropia koreana* (Vergés *et al*., 2013; Yang *et al*., 2020), a species native to the Northwest Pacific. Such insights invariably result from DNA-assisted species identification and subsequent interpretations of biogeographic patterns. Hereby widely disjunct distribution ranges are interpreted as non-natural and therefore the result of human-mediated dispersal.

Evidently, determining the non-indigenous nature of a species becomes more difficult for historic introductions. In such cases we fully rely on DNA signatures which can point toward a non-indigenous nature of the species. DNA-assisted identification of historic voucher specimens of *Codium fragile* revealed that the invasive (sub)species was already introduced into the study area as early as 1845 (Provan *et al*., 2008), which is roughly a century before phycologists realised the species was actually native to the Northwest Pacific Ocean and non-indigenous to the study area as well as several other parts of the world. In the case of *Cutleria multifida*, described from Norfolk, England as early as 1801, genetic signatures point toward an introduction of the Mediterranean Sea populations from the Northwest Pacific. The Northeast Atlantic Ocean populations, however, are genetically more diverse and well-differentiated from those in Japan and are therefore considered native (Kawai *et al*., 2016). It remains to be determined if *Cutleria multifida* is native to the Northeast Atlantic Ocean as well as the Pacific, or whether an even more complex history of historic introductions underlies this pattern.

### Discerning natural from human-mediated dispersal

Eventually, the possibility of introductions needs to be evaluated against historic and ongoing natural dispersal events. The recent observation of *Flabellia petiolata* from the south coast of England confronts researchers with exactly this question (Díaz-Tapia *et al*., 2020). Despite a long tradition of seaweed studies and regular surveys, *F. petiolata* was never recorded from the British Isles prior to 2013. The closest populations of the species are found in the Mediterranean Sea and the Canary Islands. A recent introduction would be the most obvious explanation. However, the English populations of *F. petiolata* could also be interpreted as a relic of a formerly more widespread Atlantic Ocean distribution. The species’ range might have been continuous during warmer periods in the Holocene, but persisted in the Northeast Atlantic Ocean in a handful of refugia during colder periods. Afterall, several native species, e.g. *Cladophora battersii, Codium bursa* and *Halopithys incurva* display similar distribution patterns (Maggs & Hommersand, 1993; Brodie *et al*., 2007b).

Quaternary climatic cycling probably also facilitated dispersal of temperate species across the tropical Atlantic Ocean connecting southern Africa with Europe. The presence of *Schizymenia apoda* in the Azores, the British Isles and Namibia may have resulted from natural amphi-equatorial dispersal events in recent geological times, but also a human-mediated introduction in the Atlantic Ocean cannot be ruled out given the presence of *S. apoda* in Australia and China (Gabriel *et al*., 2019; Gunnarsson *et al*., 2020). Natural dispersal events from the Northeast Pacific Ocean to Northern Europe through the Bering Strait may be difficult to discern from introductions (Lindstrom, 2001; Bringloe & Saunders, 2019). For instance, *Schizymenia jonssonii*, a species recently described from Iceland, may have colonised the northern Atlantic Ocean naturally via the Bering Strait but it is equally possible the species is a relatively recent introduction (Gunnarsson *et al*., 2020).

Population-level sampling and the application of genetic markers with sufficient intraspecific resolution (e.g. fast evolving spacer regions, microsatellite markers or SNP data) have the potential to shed light on natural versus human-mediated dispersal events, and more generally to help in reconstructing introduction history (Viard & Comtet, 2015), but are rarely used in studies of putative seaweed introductions. Notable exceptions include the invasive history of *Fucus* species, *Sargassum muticum* and *Gracilaria vermiculophylla*.

Coyer *et al*. (2011) demonstrated a North Pacific origin of *Fucus distichus* followed by at least two separate colonisation events of the North Atlantic Ocean prior to the last glacial maximum, which makes this species native to Europe. However, the taxon, having a predominantly northern distribution, was accidentally introduced in the Oslofjord followed by further expansion in the Kattegat region as a result of an introduction event in the late 19^th^ century (Coyer *et al*., 2002, as *F. evanescens*, currently regarded as a subspecies of *F. distichus*). *Fucus serratus* was also exported from mainland Europe to Atlantic North America, Iceland and the Faroes (Coyer *et al*., 2006; Brawley *et al*., 2009). Discharging of ballast stones in destination harbours is considered the prime source of introductions in Atlantic North America and Iceland in the 19^th^ century. The *F. serratus* population in the Faroes is of more recent origin (late 20^th^ century) and was most likely introduced from Iceland (Coyer *et al*., 2006). Many marine benthic organisms including seaweeds but also many invertebrates display amphi-Atlantic distribution (Haydar, 2012). For such disjunct distributions, distinguishing scenarios of post-glacial relicts or natural long-distance dispersal from human-assisted dispersal (and introduction) is challenging and most often requires a combination of life-history traits assessment and high resolution molecular markers. The power of genome-wide genetic variation was demonstrated for *Sargassum muticum* (Le Cam *et al*., 2020); whereas microsatellite markers failed to reveal any genetic variation in the invaded range of the species, a panel of single-nucleotide polymorphisms (SNPs) obtained from ddRAD sequencing confirmed a secondary introduction to the Northeast Atlantic Ocean from the Northeast Pacific Ocean, but also revealed two additional cryptic introductions to Europe. Similarly, Krueger-Hadfield *et al*. (2017) identified the areas in the native region that most likely contributed to the European invasions of the red alga *Gracilaria vermiculophylla*. Subsequent work used SNPs to refine the origins and understand evolution during invasion (Flanagan *et al*., 2021).

In Macaronesia, several confounding factors further complicate the interpretation of the non-indigenous nature of species. The geographical location of Macaronesia, bordering the tropical Atlantic Ocean, contributes significantly to this difficulty in interpretation. Several tropical and subtropical taxa are, probably erroneously, attributed a pantropical distribution, which not only contributes to high taxonomic uncertainty, but the latter also translates into biogeographic uncertainty. In addition, it is not always evident to preclude natural dispersal to explain the presence of particular species. *Halimeda incrassata*, a species that naturally occurs in the tropical western Atlantic Ocean (Verbruggen *et al*., 2006), was reported from Porto Santo, Madeira, by Wirtz & Kaufmann (2005) and more recently from the Balearic Islands by Alós et al. (2016). Further surveys indicated the species is also present in the Canary Islands (Sangil *et al*., 2018) and Azores (Costa *et al*., 2017). Even though in the Mediterranean Sea the species displays typical invasive behaviour, the involvement of human activities in its establishment is not clear. Another species from tropical western Atlantic Ocean, *Caulerpa ashmeadii*, was recently reported from Porto Santo, Madeira, and may represent a similar case of natural range expansion across the Atlantic (Ribeiro *et al*., 2023). Amphi-Atlantic Ocean distributions have been confirmed using molecular markers for several seaweed taxa (*Cladophoropsis membranacea*, Leliaert *et al*., 2009; *Laurenciella marilzae*, Cassano *et al*., 2012; *Laurencia catarinensis*, Machin-Sanchez *et al*., 2012; Tronholm *et al*., 2012; *Vertebrata foetidissima*, Díaz-Tapia *et al*., 2013; *Dictyota* spp., Tronholm *et al*., 2013; *Caulerpa prolifera*, Varela-Álvarez *et al*., 2015; *Laminaria digitata*, Neiva *et al*., 2020; *Lobophora* spp., Vieira *et al*., 2020). In these examples presumed natural distribution ranges have not been challenged.

Also of note is that Macaronesia covers a large geographic area, encompassing several biologically diverse archipelagos. Several taxa that have long been reported from the Canary Islands and Madeira and are considered native in those areas, have recently been reported from the Azores. The geographic position of the Azores and the actual oceanographic current circulation in the North Atlantic Ocean would not seem favourable for natural range expansions from the Canary Islands and Madeira. In some cases, initial reports from anthropogenic habitats, such as harbour environments (e.g. *Caulerpa webbiana*), favour the hypothesis of an anthropogenic factor in the range expansion, although evidence is lacking for several other species which are presumed non-indigenous (e.g. *Halimeda incrassata*, *Xiphosiphonia pinnulata*, *Hypoglossum heterocystideum*).

A final category of uncertainty in natural vs. human-mediated dispersal mechanisms concerns those non-indigenous species which have been displaced within the study area. The biogeographic history of the Mediterranean Sea biota is closely intertwined with the Northeast Atlantic Ocean to which it is connected by the narrow Strait of Gibraltar (Bianchi & Morri, 2003; Patarnello *et al*., 2007; Le Gall *et al*., 2021), which results in a subset of species being shared between both regions. However, several Northeast Atlantic Ocean species have been recently introduced into the Mediterranean Sea, often in lagoons with extensive aquaculture facilities, e.g. *Ascophyllum nodosum*, *Chorda filum*, *Fucus spiralis* and *Grateloupia minima* (Petrocelli *et al*., 2013). In some cases however, patterns become more complex, for example, when native and non-indigenous populations co-occur as is the case for *Chondria coerulescens*, *Vertebrata fucoides* and possibly also *Ganonema farinosum* (Verlaque *et al*., 2015). It is worth remembering that many Mediterranean Sea species naturally dispersed from the Atlantic Ocean after the Zanclean flood which occurred after the Messinian salinity crisis about 5.33 myr (Blondel *et al*., 2010).

### Species of unresolved origin

In several cases the non-indigenous nature of certain seaweeds remains unresolved. Some recently described species in the study area, such as *Polysiphonia radiata* and *P. delicata*, are mainly known from marinas and are probably non-indigenous but their origin remains unknown because it is likely that they remained undescribed in their native area (Díaz-Tapia *et al*., 2017). Morphological similarity of putative non-indigenous species to native species can also complicate interpretation of non-indigenous patterns as demonstrated by *Anotrichium furcellatum*. The latter was originally described from Naples, but considered non-indigenous in the Northeast Atlantic Ocean. The Mediterranean Sea populations, however, may have been largely replaced by a cryptic introduction of *A. okamurae*, originally from the Northwest Pacific Ocean (Verlaque *et al*., 2015). The status of *A. furcellatum* and *A. okamurae* has not yet been tested with molecular data.

Similar arguments could be made for species that are considered native in the study area. In the case of *Lobophora delicata*, which is not considered as non-indigenous, a lack of baseline data makes it difficult to be conclusive on its status as a native species. As pointed out by Vieira *et al*. (2019), the first records of *Lobophora* in the Mediterranean Sea date back to 1955 (Edelstein, 1960), which is surprising for a distinctive seaweed genus which can be easily found in many places growing at a depth of 0.5 m. In contrast, other genera of Dictyotales were invariably reported from the Mediterranean Sea in the 18th or early 19th century. Has *L. delicata* been overlooked or does the late discovery of the species correspond to a more recent introduction? Without proper baseline data, e.g. herbarium records, this is difficult to test, and if the species does not display typical invasive behaviour its native status may simply never come into question.

A puzzling case is presented by several taxa with clear Indo-Pacific affinities which appeared in the Mediterranean Sea prior to the opening of the Suez Canal in 1869, e.g. *Acanthophora nayadiformis, Asparagopsis taxiformis* and *Ganonema farinosum.* For instance, *Asparagopsis taxiformis* was first described from Alexandria in the Mediterranean Sea as *Fucus taxiformis* Delile (1813), and thus reported as a native species. However, further molecular work revealed that this accepted species was made of five distinct lineages, possibly corresponding to two cryptic species (Ní Chualáin *et al*., 2004; Andreakis *et al*., 2007; Dijoux *et al*., 2014), one of them presumably present in the Mediterranean Sea prior to the opening of the Suez Canal, and one more recently introduced. Similar complexity was revealed for the closely related species *Asparagopsis armata*, supposedly introduced in the study area, for which novel sampling in the South Pacific Ocean showed the existence of two highly divergent clades, presumably corresponding to two cryptic species, one of them distributed in Europe, South Africa and Tasmania, and one restricted (so far) to Western Australia, New Zealand and Tasmania (Dijoux *et al*., 2014). Such cases highlight the difficulty in establishing whether a species is non-indigenous in the absence of large sampling encompassing the global distribution of the targeted presumably non-indigenous species.

### The spatial patterns and origins of non-indigenous seaweeds

Analysis of the distribution of non-indigenous seaweeds in the study area reveals clear patterns in richness and the number of species shared among regions. The large-scale spatial patterns are discussed below in a context of dispersal vectors that determine spread and establishment of non-indigenous species.

The Eastern Mediterranean Sea is home to the highest number of non-indigenous seaweeds (77 species), followed by the Western Mediterranean Sea (47 species) and Lusitania (45 species). Macaronesia and Northern Europe harbour somewhat lower numbers (36 and 40 species, respectively). These numbers refer to high-confidence non-indigenous species only. Adding cryptogenic and data deficient species further underscores the higher number of non-indigenous species in the Eastern Mediterranean Sea. In the latter region an extra 47 species are flagged as cryptogenic or data deficient, which is considerably higher compared to the other regions which typically host 20 or less cryptogenic and data deficient species. In all regions, most of the non-indigenous seaweeds belong to Rhodophyta (between 63-76% of the species non-indigenous in each region), followed by brown seaweeds (18-22%), while green seaweeds contribute to 8-18% of the species recorded (Fig. 5).

**FIGURE 5.**
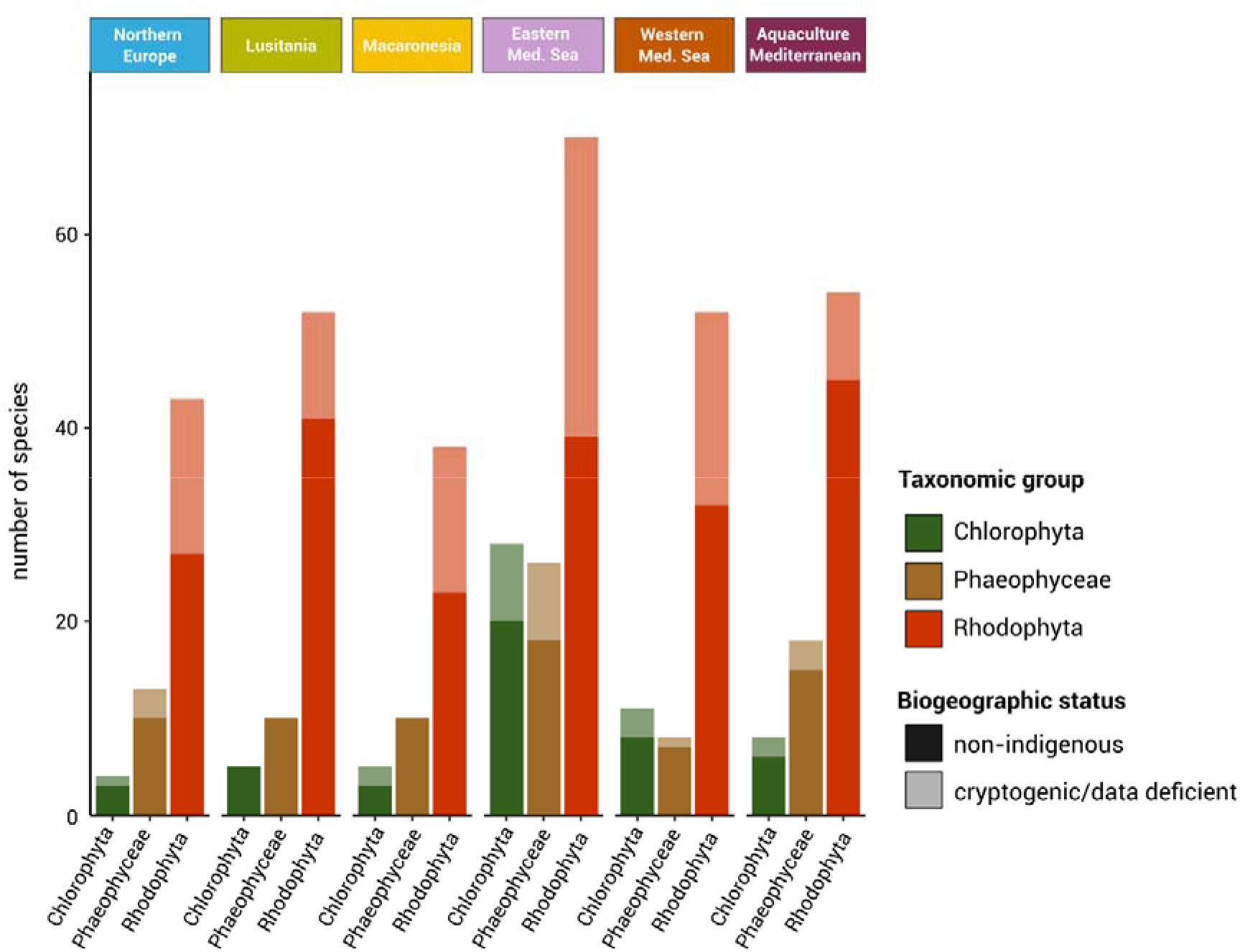
Number of non-indigenous green (Chlorophyta), brown (Phaeophyceae) and red (Rhodophyta) non-indigenous species in Northern Europe, Lusitania, Macaronesia, the Eastern and Western Mediterranean Sea, and aquaculture sites in the Mediterranean Sea. Dark shaded colours represent numbers of high-confidence non-indigenous species; light shaded colours represent cryptogenic and data deficient species.

We did not detect a significant correlation between the number of non-indigenous species and the number of records (Pearson correlation = -0.18, p-value = 0.74), which indicates that differences of non-indigenous species between regions are not a mere artefact of sampling effort. The number of non-indigenous species also does not scale with the length of the coastline (Pearson correlation = 0.10, p-value = 0.88). As will be argued below, the number of non-indigenous species in a given region and the fraction of species shared between regions is a complex function including the efficiency of primary and secondary dispersal vectors combined with abiotic (and potentially biotic) ecological factors that determine the establishment of non-indigenous species in the recipient ecosystems (reviewed in Maitner *et al*., 2021).

The Mediterranean Sea and the Northeast Atlantic Ocean share 45 high-confidence non-indigenous species, while Macaronesia shares roughly an equal number of non-indigenous species with the Mediterranean Sea (24 species) and the Northeast Atlantic Ocean (21 species) (Fig. 6A). Within the Northeast Atlantic Ocean, Macaronesia and Lusitania share 22 high-confidence non-indigenous species, while Macaronesia and Northern Europe share none other than the 14 non-indigenous species present in all three Northeast Atlantic regions (Fig. 6B). A relatively low number, 18 high-confidence non-indigenous species out of 140, are shared between the Northeast Atlantic Ocean, Macaronesia and the Mediterranean Sea (Fig. 6A). The broad distribution of these non-indigenous species is noteworthy for it reflects a very wide amplitude in abiotic and biotic parameters. At least nine of these widely distributed non-indigenous species (*Antithamnion hubbsii/nipponicum, Antithamnionella spirographidis, Asparagopsis armata, Bonnemaisonia hamifera, Codium fragile* subsp. *fragile, Colpomenia peregrina, Dictyota cyanoloma, Grateloupia turuturu, Scytosiphon dotyi*) are reported from all five regions. The remaining species have a more restricted distribution range, being only present in the three central regions Macaronesia, Lusitania and the Western Mediterranean Sea (*Phycocalidia suborbiculata, Spongoclonium caribaeum, Symphyocladia marchantioides*), or being absent from either of the two peripheral regions, i.e. the Eastern Mediterranean Sea (*Antithamnionella ternifolia, Sargassum muticum*) or Northern Europe (*Asparagopsis taxiformis, Rugulopteryx okamurae, Pachymeniopsis gargiuloi*).

**FIGURE 6.**
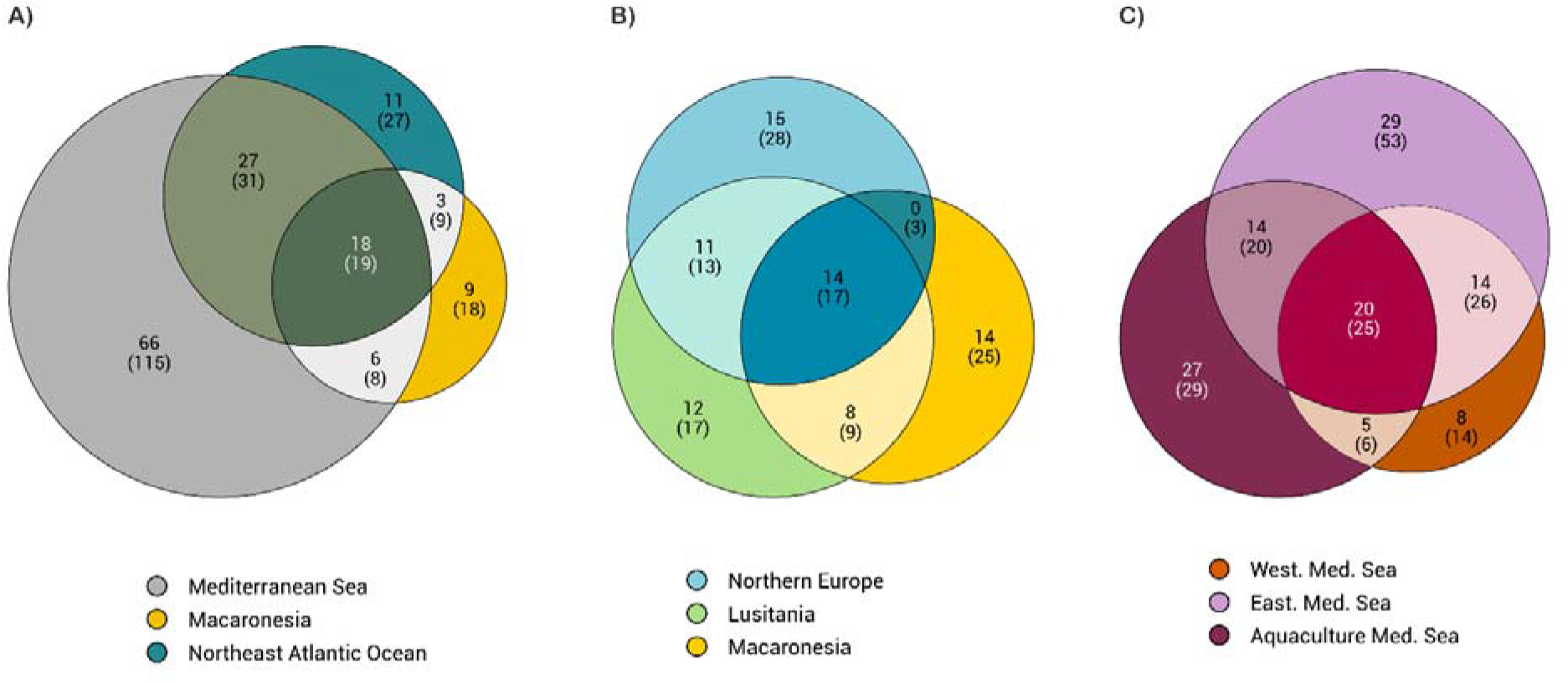
Number of non-indigenous species shared among regions. A) Mediterranean Sea, Macaronesia, and Northeast Atlantic Ocean; B) Northern Europe, Lusitania, and Macaronesia; C) Western Mediterranean Sea, Eastern Mediterranean Sea, and aquaculture sites in the Mediterranean Sea. Numbers display the high-confidence non-indigenous species (excluding cryptogenic and data deficient species), with total number of non-indigenous species including cryptogenic and data deficient species in brackets.

The number of high-confidence non-indigenous species currently only found in the Mediterranean Sea (66 species) is striking. Including cryptogenic or data deficient species brings this number to 115. However, there is considerable differentiation of non-indigenous species between the Western and Eastern Mediterranean Sea. Only 34 non-indigenous species are shared between both regions, representing 38% of the total non-indigenous species diversity in the Mediterranean Sea (Fig. 6C). In addition, the fraction of non-indigenous species unique to the Eastern Mediterranean Sea (27 species) is considerably larger than for the Western Mediterranean Sea (8 species) (Fig. 6C). The latter pattern is largely the result of dispersal of warm-adapted species from the Red Sea and by extension the Indo-Pacific Ocean via the Suez Canal. At present, only a fraction of these have spread to the Western Mediterranean Sea resulting in a higher diversity of non-indigenous species in the Eastern Mediterranean Sea.

The non-indigenous species reported from the Thau Lagoon in France, and the Mar Piccolo and Venice Lagoon in Italy are quite distinct compared to those of surrounding Mediterranean waters. A combination of anthropogenic disturbances and intense aquaculture activities (trade and exchanges), more specifically import of shellfish, has resulted in a very high diversity of non-indigenous species in these lagoon systems (66 species), comparable to that of the surrounding Mediterranean Sea locations despite their much smaller area. The non-indigenous species in Mediterranean lagoons have, moreover, more affinities with the Atlantic Ocean than with the Mediterranean Sea, likely to be due to exchanges between shellfish production areas (see below). Nearly half of the non-indigenous species encountered in Mediterranean aquaculture lagoons are not (yet) reported from surrounding Mediterranean water, while 36 are shared with the Northeast Atlantic Ocean. Of these 36 non-indigenous species, 13 have only been recorded in aquaculture lagoons within the Mediterranean Sea. These mainly include species with a Northwest Pacific Ocean origin, e.g. *Dasysiphonia japonica*, *Neopyropia yezoensis* and *Nitophyllum stellatocorticatum*. Differences in the abiotic physico-chemical environment between Mediterranean lagoons and surrounding coastlines probably underlie the failure of these species to spread widely in the Mediterranean Sea. *Rugulopteryx okamurae*, however, presents a striking counterexample of this trend. The species was collected in the Thau Lagoon for the first time in 2002 (Verlaque *et al*., 2009). Initially *R. okamurae* appeared to be rather non-invasive, but in 2015 it was reported from the Strait of Gibraltar (Ocaña *et al*., 2016; El Aamri *et al*., 2018; García-Gómez *et al*., 2020), where the species forms dense stands rapidly overgrowing most native seaweed species. More recently the same alarming invasive behaviour of *R. okamurae* has been noted in the Marseille area (Ruitton *et al*., 2021) as well as southwest Portugal (Liulea *et al*., 2023) and Macaronesia (Faria *et al*., 2022).

A Northwest Pacific origin of the largest part of the non-indigenous species present in the Mediterranean lagoons and Northern Europe is well established (Fig. 7, Table 1) (Boudouresque *et al*., 2010). Regular monitoring and surveys demonstrated that many of those species had been first accidentally introduced in the Mediterranean lagoons (notably the Thau lagoon) and were then transported to the Northeast Atlantic Ocean hitchhiking with oyster transfers (Mineur *et al*., 2007a). *Undaria pinnatifida* presents a notable exception to this pattern. Following its accidental introduction in the Thau lagoon, the species was deliberately introduced in Brittany for aquaculture purposes (Floc’h *et al*., 1991) from which it rapidly spread and established itself as one of the dominant non-indigenous species in artificial as well as natural habitats in the Northeast Atlantic Ocean (Voisin *et al*., 2005; Guzinski *et al*., 2018). Note that commercial transfers of oysters between the Mediterranean Sea and the Atlantic coasts of France is still fully allowed under French and European regulations, which results in a quasi-continuous series of secondary introduction events. Aquaculture- and fisheries-associated transport (e.g. nets, packing material) between the Atlantic Ocean and Mediterranean Sea is also undoubtedly responsible for the introduction of a range of native Atlantic Ocean species to the Mediterranean lagoons (e.g. *Ascophyllum nodosum, Chorda filum, Grateloupia minima, Fucus spiralis* and *Vertebrata fucoides*).

**FIGURE 7.**
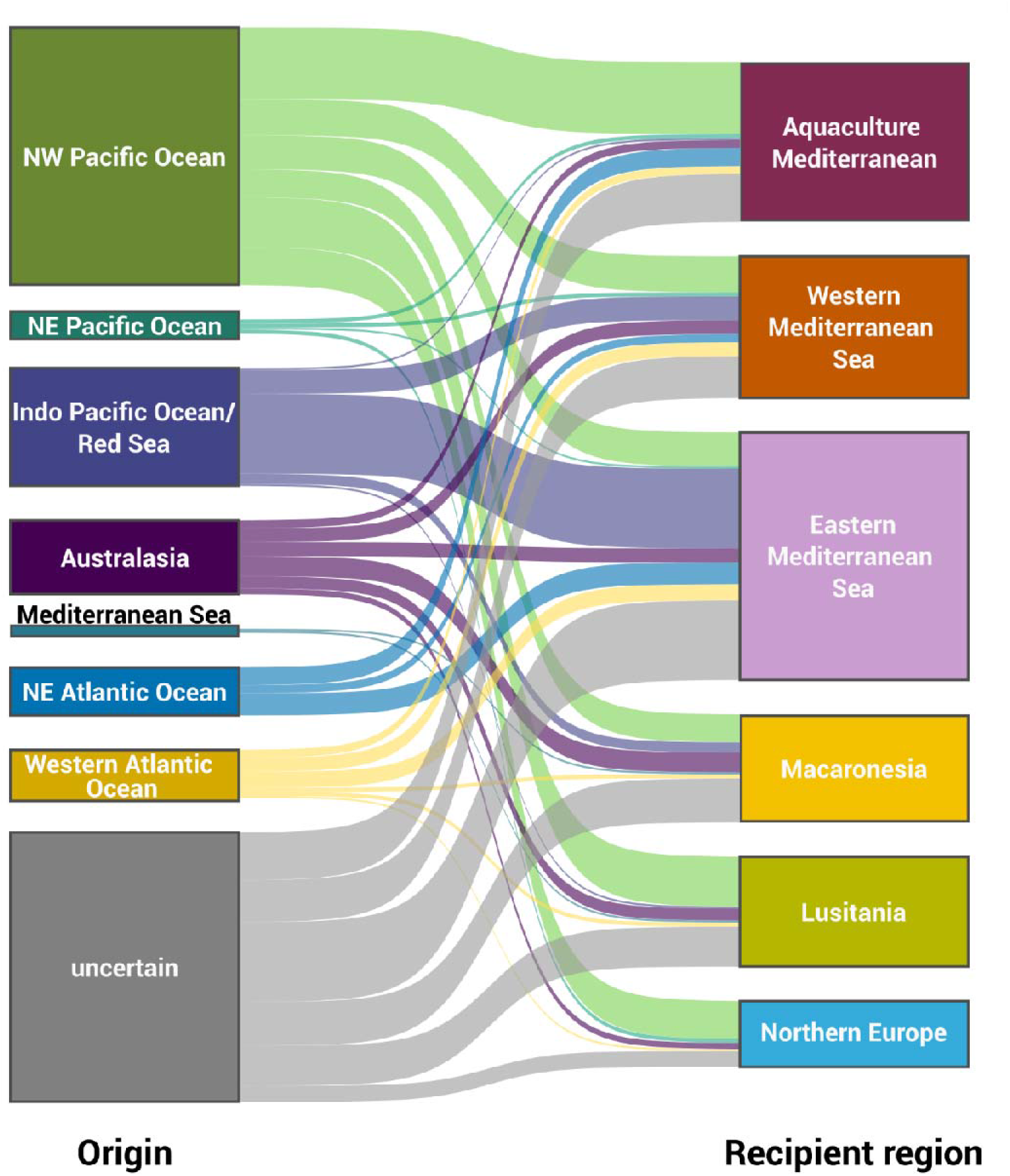
The origin of non-indigenous species. For each of the recipient study regions, the Sankey diagram displays what proportion of non-indigenous species have likely originated from a specific area (Northwest Pacific Ocean, Northeast Pacific Ocean, Indo Pacific Ocean/Red Sea, Australasia, Mediterranean Sea, Northeast Atlantic Ocean, Western Atlantic Ocean, or uncertain). The width of the flow arrows is proportional to the number of non-indigenous species (including non-indigenous, cryptogenic and data deficient species).

The Mediterranean Sea, Macaronesia and Lusitania are also home to 35 high-confidence non-indigenous species with presumed or established Australasian origin (Fig. 7). Species with Australasian origin are much less represented in Northern Europe, reflecting the warm temperate nature of these species. The introduction vectors for this category of species remain, however, largely elusive. For *Acrothamnion preissii* and *Womersleyella setacea* ship traffic has been suggested as vector based on their first observation close to a major harbour (Livorno, Italy), but accidental release from scientific laboratories and public or private aquaria is also a possibility. As with the notorious case of *Caulerpa taxifolia* (Verlaque *et al*., 2015), aquarium releases have likely resulted in the introduction of other seaweeds, mainly in the Mediterranean Sea (Vranken *et al*., 2018).

Despite the abovementioned clear categories of non-indigenous species and associated pathways, for circa one third of the species there is considerable uncertainty regarding the area of origin and the potential dispersal vectors. Complicating identification of native range and vectors even further, population-level molecular studies on several non-indigenous species have unveiled multiple independent introductions possibly involving different vectors (McIvor *et al*., 2001; Voisin *et al*., 2005; Geoffroy *et al*., 2016; Le Cam *et al*., 2020).

### Introduction hotspots

Of the 140 high-confidence non-indigenous species, 65% have been reported for the first time in the Mediterranean Sea, 26% in the Northeast Atlantic Ocean and 9% in Macaronesia. The distributions of the first record of each species in the study area underscore the importance of aquaculture for introductions of seaweeds. The Thau lagoon, with 30 reports of first introductions in the study area, is one of the major introduction hotspots. In total 58 species, constituting 32% of the total seaweed diversity or 48-99% of the biomass, have become established in this coastal lagoon (Boudouresque *et al*., 2010). The Thau lagoon is an important centre of oyster cultivation in the Mediterranean Sea. However, the oyster farmers rely on the import of oyster spat produced in other regions because the lagoon is not particularly suitable for oyster reproduction. Since 1977, only spat of Pacific oysters spat produced in the Atlantic Ocean is allowed to be laid in the French Mediterranean lagoons (Verlaque *et al*., 2007). However, it is likely that some non-official imports from outside of Europe occur, as reported by Verlaque (1996). These continuous transfers across different biogeographic regions result in astonishingly high numbers of non-indigenous species. A low native diversity due to the low occurrence of natural hard substrata in lagoons and relatively recent construction of hard substrata for aquaculture purposes, concomitant with transfers of livestock which seed the new substrata, makes these lagoons hotspots for non-indigenous species establishment (Mineur *et al*., 2015).

The Southeast Mediterranean Sea constitutes another introduction hotspot, which accounts for 24 first reports and a total of 32 non-indigenous species. The inauguration of the Suez Canal in 1869 resulted in an open connection between the Northern Red Sea and the Eastern Mediterranean Sea. As a result, more than 500 marine species are believed to have invaded the Mediterranean Sea through the Suez Canal, so-called Lessepsian migrants (Zenetos *et al*., 2010; Zenetos *et al*., 2012; Galil *et al*., 2021). With respect to non-indigenous seaweeds, many species were first reported in a series of papers by the Egyptian phycologist Anwar Aleem (Aleem, 1948; Aleem, 1950; Aleem, 1951; Aleem, 1993). Recent efforts by Greek, Lebanese, Israeli and Turkish phycologists have expanded the list of Lessepsian seaweeds considerably (e.g. Tsiamis, 2012; Hoffman, 2013; Bitar *et al*., 2017; Israel & Einav, 2017; Çinar *et al*., 2021; Galil *et al*., 2021). Nevertheless, a paucity of historic baseline data makes it often difficult to establish the Lessepsian origin of many species or to point to the exact date of introduction. As outlined above, records of species with clear Indo-Pacific affinities which predate the opening of the Suez Canal (e.g. *Ganonema farinosum* and *Acanthophora nayadiformis*) still puzzle phycologists. In addition, the identities of many species reported for the first time by Aleem and others (e.g. *Gracilaria arcuata, G. disticha, Hypnea flagelliformis*, *Solieria dura, Spatoglossum variabile*) have never been confirmed using molecular markers and are highly uncertain. In general, a detailed understanding of past and contemporary temporal dynamics of seaweed introductions in the Eastern Mediterranean Sea remains a challenge. More than in any other region it remains difficult to link the observation of a new seaweed species with the introduction date. This uncertainty has bearing on the monitoring of migration through the Suez Canal which has been regarded as an ongoing process (Boudouresque, 1999; Por, 2012). The current construction of the new Suez Canal, doubling the capacity of the current corridor, is expected to further increase the influx of Red Sea species (Galil *et al*., 2015) and contribute to further tropicalisation of the Mediterranean Sea (Bianchi & Morri, 2003; Coll *et al*., 2010).

Compared to the two Mediterranean Sea introduction hotspots, first records appear less localised in the Northeast Atlantic Ocean. The English Channel (Brittany, southern English coast) and the Scheldt estuary (the Netherlands) are most prominent as introduction hotspots. To what extent this spatial pattern reflects the true locations of primary introductions or whether the locations of first records are biased by the distribution of preferred study areas of phycologists and research institutes is difficult to assess. There is, indeed, a strong correlation between the introduction hotspots (i.e. locations from which a high number of first records for the study area were reported) and the density map of all records of non-indigenous species, which is indicative of high monitoring activities in areas where many non-indigenous species are found. In addition, it is noteworthy that the English Channel and Scheldt estuary are important areas for oyster farming, and many non-indigenous species have been accidentally introduced with oyster transfer from the Mediterranean Sea.

### Introduction rates: temporal trends

Disentangling the factors underpinning temporal trends in the accumulation rate of non-indigenous seaweeds may improve our understanding of introductions and result in better-informed predictions of future trajectories (Seebens *et al*., 2018). Deducing temporal trends in the rate of introduction of non-indigenous species, however, assumes a correlation between the date of introduction and the moment the species was detected. Although seemingly straightforward, for several species the timespan between introduction and detection is probably considerable and unpredictable. Detection obviously depends on collecting effort, but as highlighted in previous sections, the taxonomic and biogeographic framework will also determine if a species is considered non-indigenous.

Furthermore, Costello & Solow (2003) demonstrated that an increasing rate of detection need not imply an increasing rate of introductions even when collection effort is constant. Given this complexity, reports that introduction rates have increased or decreased in specific time windows should be treated with caution.

Acknowledging this uncertainty, the detection of non-indigenous species shows two distinct phases, one prior to 1950-1970 characterised by low accumulation rates, followed by another much higher accumulation rate from then onward (Fig. 8). Irrespective of taxonomic and biogeographic uncertainties, there is little to no indication for a decline in the rate at which non-indigenous species are reported. The observation that the detection, and presumably also the introduction, of non-indigenous species has not reached saturation (Fig. 8), is in line with the observations by Seebens *et al*. (2017) for other taxonomic groups. For seaweeds, the absence of a decline in the first-record rate may point to the inefficiency of measures aimed to prevent and mitigate new introductions.

**FIGURE 8.**
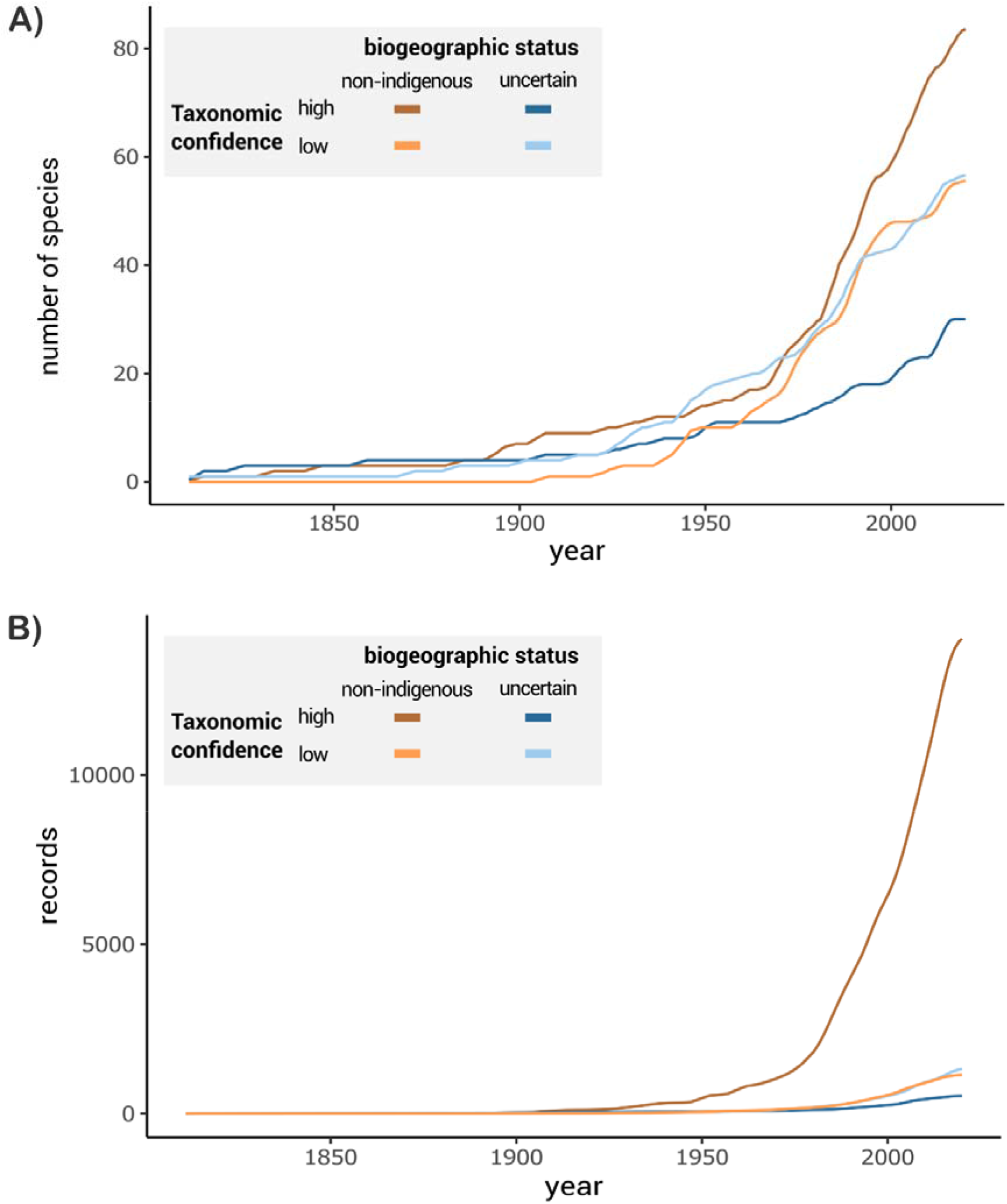
The accumulation of A) the number of records, and B) the number of non-indigenous species through time (1808-2022). Colours indicate taxonomic confidence and biogeographic status: dark brown (taxonomic confidence = high, biogeographic status = non-indigenous), light brown (taxonomic confidence = low, biogeographic status = non-indigenous), dark blue (taxonomic confidence = high, biogeographic status = uncertain), light blue (taxonomic confidence = low, biogeographic status = uncertain). All trends are displayed as moving averages over 5 years.

Alternatively, the community of phycologists involved in monitoring of non-indigenous seaweeds has become larger and more efficient at detecting incoming species, e.g. through the use of DNA-barcoding methods.

According to Mineur *et al*. (2007b) who sampled the hulls of several commercial cargo vessels, hull fouling seems to play a relatively minor role in the displacement of seaweeds across the globe. Similarly, ballast water, one of the prime sources of introductions for marine invertebrates and microalgae (Bolch & de Salas, 2007; Gollasch *et al*., 2015), is relatively unimportant with respect to introduction of seaweeds. Yet there is evidence of introduction with commercial ships not hulls or ballast water but other components such as anchors, for instance in Australia for *U. pinnatifida* (South *et al*., 2017). Leisure boats, however, likely contribute to the local spread of non-indigenous species within the study area. Their role has certainly been underestimated so far, at least for secondary introductions (Mineur *et al*., 2008). Contrary to the Eastern Mediterranean Sea where the opening of the Suez Canal resulted in an ongoing and steady influx of non-indigenous species (Galil *et al*., 2015), in the Western Mediterranean Sea a disproportionate number of non-indigenous seaweed species appears to have been introduced through import of oyster stocks (Verlaque *et al*., 2007). In the late 1960s and early 1970s, disease caused by Asian oysters importation outbreaks in the study area affecting oyster populations caused a major disruption of production (Mineur *et al*., 2015). Mitigation procedures involved massive imports in the 1970s of Pacific oyster from its native range in the Northwest Pacific Ocean, or via from the Puget Sound in the Northeast Pacific Ocean where the species is also cultivated on a large scale (Mineur *et al*., 2014). *Sargassum muticum* introduced into the study area was shown to have several origins including a primary introduction from Asia and a secondary introduction from the Northeast Pacific Ocean (Le Cam *et al*., 2020).

## Conclusion and perspectives

In conclusion, our critical synthesis of non-indigenous seaweed diversity in the Northeast Atlantic Ocean, Mediterranean Sea and Macaronesia revealed widespread taxonomic and biogeographic uncertainty. This finding negatively impacts efforts to evaluate the effectiveness of measures to reduce non-indigenous species influx, manage their risks and impacts, and devising potential control strategies. This uncertainty can be addressed through the progressive use of molecular markers, particularly standard DNA barcoding approaches, which can in most cases confirm the identification of presumed non-indigenous seaweeds (Viard & Comtet, 2015). Importantly, however, a taxonomic and biogeographic reference framework should also be established for putative indigenous regions. DNA-based identification is relatively well-developed for species with a Northwest Pacific origin. For tropical taxa such a framework lags behind, resulting in higher levels of uncertainty regarding the identity of non-indigenous species with presumed tropical origin. Such a reference framework will also be necessary for early detection of non-indigenous species and monitoring with the use of bulk sample and eDNA metabarcoding (Darling *et al*., 2017; Keck & Altermatt, 2023).

However, standard DNA barcoding may not be sufficient to interpret more complex introduction histories, such as cases where non-indigenous and indigenous populations co-occur, as suggested for several species in the Mediterranean Sea (Verlaque *et al*., 2015) or where recent human-mediated dispersal needs to be evaluated against ongoing and natural dispersal events linked to Pleistocene or Holocene climatic oscillations (Neiva *et al*., 2016). In these cases a combination of population-level sampling strategies and molecular markers that capture intraspecific diversity is needed to shed light on the number and directions of dispersal events and re-evaluate the status of taxa currently considered cryptogenic.

In parallel with DNA-barcoding efforts, historical and reliable baseline data of seaweed diversity is needed to reduce the subjective interpretation of non-indigenous species and determine more precisely their date of introduction. It is likely that the introduction of many non-indigenous species occurred significantly earlier than the time of their first detection, especially for species that have morphologically similar congeners in the study area (such as species from the genera *Dictyota*, *Gracilaria*, *Polysiphonia* and *Ulva*). Herbarium collections can serve as a crucial source of primary data to address this issue, and advances in sequencing technologies make it possible to obtain genetic data from voucher specimens tens or even hundreds of years old as demonstrated for the *Codium fragile*-complex (Provan *et al*., 2008). We anticipate that herbaria will play an increasingly important role in documenting spatio-temporal patterns of non-indigenous seaweeds, alongside large-scale digitization efforts for these collections. More precise estimates of the date of introduction will also reduce uncertainty regarding the accumulation rate of non-indigenous species. Our analyses suggest that the rate of introduction in the study area has not decreased. However, it is unclear whether this trend reflects a steady accumulation of non-indigenous species or increased and more efficient detection through monitoring or advanced identification methods. Based on reliable baselines, temporal surveys will allow to uncover the trends in non-indigenous species introduction rates. Diversifying the type of survey, from punctual surveys (e.g. Bio-Blitzes or Rapid Assessment surveys) to systematic comprehensive surveys (e.g. full inventories), including morphological-based and DNA-based assessments, such as metabarcoding (see above), is a need for effective prevention and early detection and also to monitor trends over time in new species introductions.

Our study also reveals significant differences in the geographical distribution of non-indigenous seaweed species across the study area, with only 18 species shared between the three main regions. Non-indigenous species distribution reflects both their abiotic niche and the efficiency of primary and secondary dispersal vectors. It is expected that current patterns will become increasingly homogenous with time due to various factors. First, recently introduced species may continue to expand as part of the ongoing invasion process. Secondly, evolutionary processes such as selection, genetic admixture and hybridisation may occur during the invasion process, leading to adaptation and expansion of the species in new ecological conditions. Last, further range expansions are expected for non-indigenous species with affinities for warm temperate to tropical temperatures as a result of ongoing ocean warming. This could result in an influx of warm-water adapted species into regions where conditions are currently unfavourable, while warming may also render regions unfavourable for non-indigenous species that currently thrive there. It is anticipated that under ocean warming, Eastern Mediterranean Sea non-indigenous species would likely expand their range in western direction, while species ranges in Lusitania would shift northward. The accurate estimation of the rate at which non-indigenous species expand their ranges, either as invasion fronts or more erratically by jump dispersals, rely on detailed monitoring. Such estimates can inform on dispersal vectors at various spatial scales and guide policy makers to take effective measures to prevent or limit the spread of these species.

We advocate that combining the efforts of taxonomists who provide a reliable framework on the number and likely geographic origin of non-indigenous seaweeds, together with environmental monitoring, offers the best strategy to identify species of concern, characterise their life history traits, and develop effective management strategies. Especially for the species entering the Mediterranean Sea through the Suez Canal, a combination of horizon scanning exercises, intensive monitoring and rapid-response eradication efforts at the local level may be the only tools to try and control the establishment of Lessepsian migrants. With respect to non-indigenous species that hitchhike with shellfish transport, effectively limiting imports into Europe and controlling translocations between regions in Europe should effectively reduce the rate of primary and secondary introductions (Mineur *et al*., 2014). Alternatively, immersion for shorter periods (3 seconds) at temperatures of 80–85°C is effective in killing macroalgal propagules (Mineur *et al*., 2007a). In addition, hull fouling has an important role for introductions, especially towards secondary spread of already introduced non-indigenous seaweeds (Clarke Murray *et al*., 2011). The related guidance developed in the context of the Marine Environmental Protection Committee (MEPC, 2011) is a step forward, but we stress the need for more enforceable control of this pathway. Mitigating the negative effects of non-indigenous species that have already established will likely prove to be even more difficult. Solid baseline data will allow us to detect introduction patterns and non-indigenous species range shifts in early stages and act accordingly.

Combatting the effects of non-indigenous seaweeds will require coordinated action at the European and international level to prevent the introduction of species, to quickly detect and rapidly eradicate species to prevent them from establishing, and to manage established species to minimise their ecological and economic impact (IAS Regulation (EU) 1143/2014 on invasive alien species). These measures will require significant efforts and collaboration between science, management, policy, and society. Our dataset supports these regulatory actions by providing a solid baseline on non-indigenous seaweeds. This baseline contributes to the assessment of the current situation, helps authorities to identify new introductions and monitor the status of already established species, and importantly identifies the current knowledge gaps concerning taxonomic and biogeographic uncertainties.

## Acknowledgments

The research was carried out as part of the ERANET INVASIVES project (EU FP7 SEAS-ERA/INVASIVES SD/ER/010), with infrastructure provided by EMBRC Belgium - FWO project GOH3817N and I001621N. This paper benefited from surveys and studies that FV carried out in the IDEALG (ANR-10-BTBR-04) and Interreg IVa Marinexus projects. This is publication ISEM 202X-XXX of the Institut des Sciences de l’Évolution -Montpellier. ES acknowledges EU-BiodivERsA BiodivRestore-253 (RESTORESEAS, funded by FWO and FCT). AP and EC acknowledge the support of activities from: 1) the Apulian Region through the POR PUGLIA FESR-FSE 2014/2020, Asse VI, Action 6.5; and 2) the Italian National Recovery and Resilience Plan (PNRR) funded by the Italian Ministry of University and Research, Mission 4, Component 2, “From research to business”: 1. NBFC, Investment 1.4, Project CN00000033.

## Supplementary information

Supplementary Table S1: concise description of the status of species flagged as non-indigenous in the Northeast Atlantic Ocean, the Mediterranean Sea, and Macaronesia.

The complete dataset containing all records is available at Zenodo (DOI:10.5281/zenodo.7798640).

## Author contributions

LM. van der Loos + Q. Bafort +S. Bosch: concept, data acquisition, analyses, writing; F. Leliaert + O. De Clerck: concept, data acquisition, analyses, writing. Other authors: data acquisition, writing.

